# Defining the contribution of microRNA-specific slicing Argonautes in animals

**DOI:** 10.1101/2023.01.19.524781

**Authors:** Anisha Pal, Vaishnav Vasudevan, François Houle, Michael Lantin, Katherine A. Maniates, Miguel Quévillon Huberdeau, Allison L. Abbott, Martin J. Simard

## Abstract

microRNAs regulate gene expression through interaction with an Argonaute protein family member. While some members of this protein family retain an enzymatic activity capable of cleaving RNA molecules complementary to Argonaute-bound small RNAs, the role of the slicing activity in the canonical microRNA pathway is still unclear in animals. To address the importance of slicing Argonautes in animals, we created *Caenorhabditis elegans* strains, carrying catalytically dead endogenous ALG-1 and ALG-2, the only two slicing Argonautes essential for the miRNA pathway in this animal model. We observe that the loss of ALG-1 and ALG-2 slicing activity affects overall animal fitness and causes phenotypes, reminiscent of miRNA defects, only when grown and maintained at restrictive temperature. Furthermore, the analysis of global miRNA expression shows that the catalytic activity of ALG-1 and ALG-2 differentially regulate the level of specific subsets of miRNAs in young adults. We also demonstrate that altering the slicing activity of those miRNA-specific Argonautes does not result in any defect in the production of canonical miRNAs. Together, these data support that the slicing activity of miRNA-specific Argonautes function to maintain the levels of a set of miRNAs for optimal viability and fitness in animals particularly exposed to specific growing conditions.

## INTRODUCTION

The microRNAs (miRNA) are ∼21 nucleotides (nt) long non-coding RNAs involved in various biological processes in animals, plants, and fungi, including proliferation and differentiation, development, circadian rhythm, and stress response(1). Typically, miRNAs bind to the complementary sequence found in the 3’ untranslated region (UTR) of target mRNA, thus facilitating post-transcriptional gene silencing (2, 3). The production of miRNA involves molecular steps that occur in the cell’s nucleus and cytoplasm, respectively. First, the primary miRNAs (pri-miRNA) are transcribed by RNA polymerase II. Following this, the microprocessor complex (Drosha-DGCR8) binds to and cleaves the pri-miRNA in the nucleus to generate the precursor miRNA (pre-miRNA) (4). The pre-miRNA is then exported into the cytoplasm, wherein it is cleaved by Dicer to produce small duplex RNA molecules, comprised of the mature miRNA guide strand and the passenger miRNA star strand. An Argonaute protein subsequently binds the mature guide strand, while the passenger strand (star miRNA) is discarded or not utilized (5–7). Following the loading of the mature miRNA, the bound Argonaute protein associates with other factors, such as TNRC6/GW182 protein, to assemble the miRNA-induced silencing complex (miRISC). The miRISC binds to the complementary 3’UTR of the target mRNA and recruits various factors to cause post-transcriptional gene silencing (8). In animals, the miRNA pathway is involved in translational repression and the degradation of the target through the recruitment of the deadenylase, CCR4/NOT complex, by TNRC6/GW182 (9). Recently, it has been shown that miRNAs can modulate gene expression in a TNRC6/GW182 independent manner in particular conditions, such as animal embryogenesis (10). Interestingly, along with blocking translation, miRISC binding can stabilize the targeted mRNAs in the germline by sequestering them at a specific subcellular location by miRISC-mediated interaction with GLH-1/Vasa (11). Therefore, interactors, besides the miRNA molecule and Argonaute, in *cis* and *trans* of the miRISC, can modulate its gene regulatory mode of action (12).

The Argonaute protein family represents the functional key player of all small non-coding RNA-mediated gene silencing pathways found across archaea to eukarya. Phylogenetically, there are three major classes of Argonautes: the Argonautes, the PIWI proteins and a set of worm-specific Argonautes involved in different small RNA-specific gene regulatory processes (13). The miRNA-mediated gene silencing involves members of the Argonaute clade, encompassing Ago1 to 4 in mammals and ALG-1, ALG-2, and ALG-5 in the nematode *Caenorhabditis elegans* (14, 15). The Argonaute proteins contain four major domains: The N-terminus, PAZ (Piwi/Argonaute/Zwille), Mid and PIWI (P-element induced wimpy testis) (16, 17). While the PAZ and Mid domains are required to bind the 3’ and 5’ end of small RNAs, respectively, the C-terminus PIWI domain can act as an endonuclease that cleaves the RNA backbone’s phosphodiester bond. This enzymatic or slicing activity requires four specific (DEDH) residues (known as catalytic tetrad) within the PIWI domain, to bind divalent cation (i.e. Mg+2, Mn+2) and cleave target mRNA, if there is near perfect or perfect complementarity with the bound small RNA molecule (14, 18).

The Argonaute slicing activity is essential for siRNA-guided target cleavage and piRNA-mediated genome integrity maintenance in the animal germline (19–25). In contrast to plants, the absence of the perfect complementarity between miRNAs and mRNA targets, prevents their cleavage in animals. Many Argonautes that bind miRNAs in animals have retained the residues essential for target cleavage (26). Previous reports have suggested the limited contributions of the slicing activity of Argonautes in the miRNA pathway. For instance, slicing human Ago2 has been demonstrated to be engaged in the production of specific functional miRNAs in cells. In vertebrates, Ago2-mediated cleavage activity produces two specific miRNAs, miR-451 and miR-486, essential for erythroid development (27–30). In the case of miR-451, its maturation creates a short hairpin structure, which bypasses Dicer and directly binds Ago2 to be processed into mature miRNA (28–30). On the other hand, miR-486 necessitates Dicer for its maturation but requires slicing Ago2 to eliminate the star strand from the miRNA duplex (27). While those studies highlight the contribution of slicing Argonaute in producing specific miRNAs, a few limitations in those approaches prevent our understanding of the broader impact of the slicing activity of Argonautes in the miRNA pathways in animals. First, Ago2 slicing activity is essential for another small RNA gene silencing pathway that uses siRNA to cleave mRNA targets (31, 32). Next, it has been recently reported that Ago3, another member of the Argonaute family implicated in the miRNA pathway, also possesses slicing activity (33). Therefore, decoupling the Argonaute slicing activity for siRNA and miRNA production and understanding its contribution to miRNA function has remained a challenge in animals.

Here, we take advantage of the nematode *C. elegans* as a model to elucidate the contribution of the slicing activity of Argonautes in the animal miRNA pathway. *C. elegans* has three Argonautes (ALG-1, ALG-2 and ALG-5), exclusively implicated in the miRNA pathway and phylogenetically related to all four human Argonautes (34–36). However, of the three, only ALG-1 and ALG-2 carry the four residues essential for slicing activity (Figure 1A) and are catalytically active (37). Therefore, we created animals, deprived of ALG-1 and ALG-2 slicing activity and studied their contributions to the miRNA pathway. We found that their catalytic activities considerably impacted animal fitness and different developmental pathways, controlled by miRNAs, when maintained in stressful conditions. We also observed that ALG-1 and ALG-2 slicing activity regulate the abundance of specific miRNAs without causing any defect in their production.

**Figure 1.**
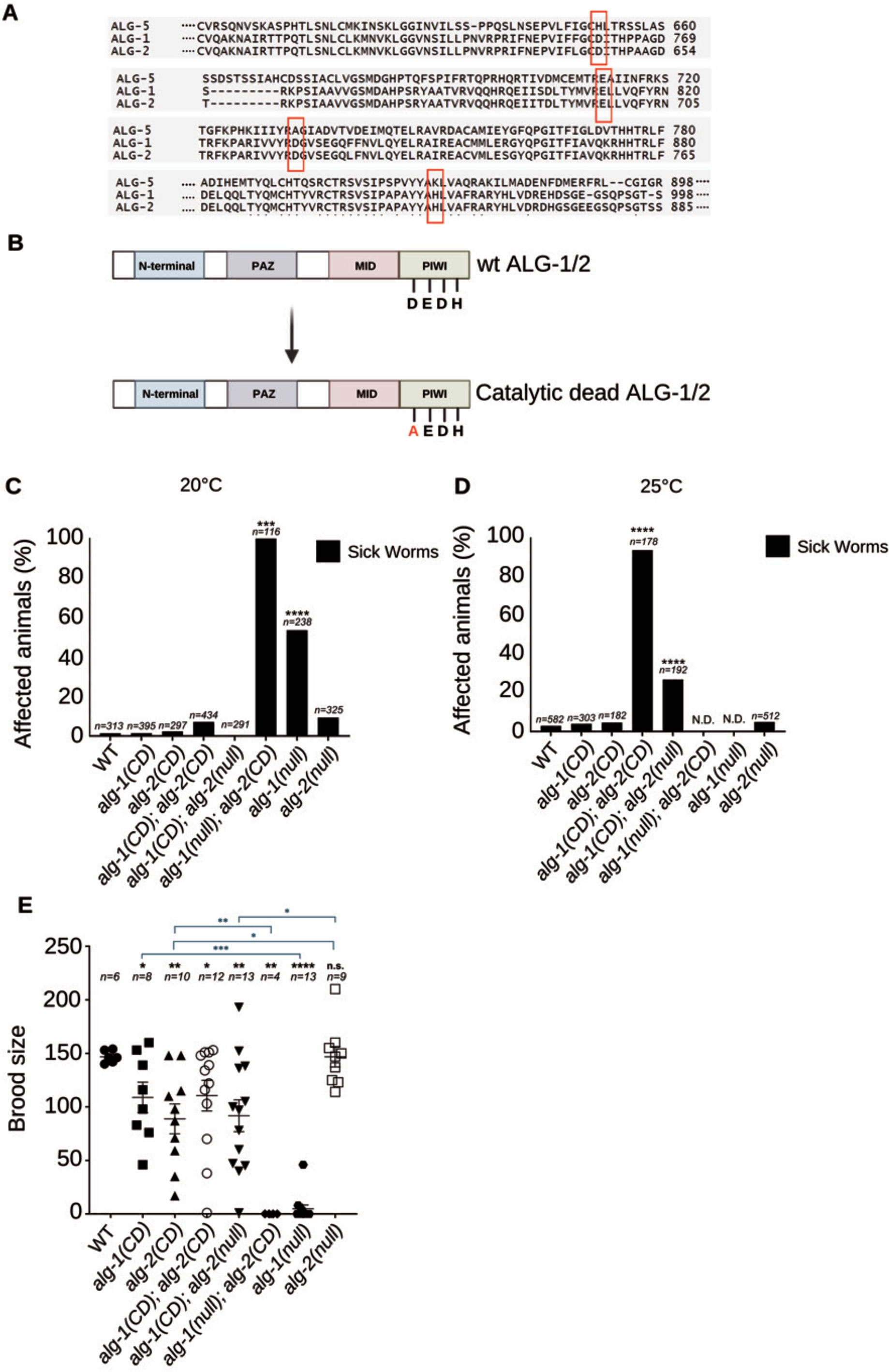
Catalytically dead ALG-1 and ALG-2 affect animal fitness. (A) The alignment of ALG-1/2/5, done on CLUSTAL O (1.2.4) showing the presence and absence of catalytic tetrad (the four amino acids DEDH are identified by the red rectangles). **(B)** Schematic representation of the four Argonaute domains (N-terminal, PAZ, MID and PIWI), the catalytic tetrad of ALG-1 and ALG-2 and generated catalytic dead endogenous Argonaute (ALG-1/2) mutants by CRISPR-Cas9 system. **(C, D)** The worm fitness was observed under normal table top microscope at two different temperatures: 20°C (panel **C**) and 25°C (panel **D**). The percentage of sick (represented by pleiotropic phenotypes such as developmental arrest, protruding vulva, egg laying defect) worms is indicated. The number of animals scored (*n*) is indicated. *p-value* was obtained by two tailed Fisher’s exact test (\**p*<0.05, \*\**p*<0.01, \*\*\**p*<0.001, \*\*\*\**p*<0.0001). **(E)** Total hatched larvae were counted for P0 mothers of each genotype. The mean brood size as well as S.E.M. per genotype are indicated. The number of animals scored (*n*) is indicated. *p-value* was counted using Kolmogorov test (\**p*<0.05, \**\*p*<0.01, \*\*\**p*<0.001, \*\*\*\**p*<0.0001) to include the extreme data points.

## MATERIAL AND METHODS

### Worm culture and Phenotypic studies

All *C. elegans* strains were cultured and handled using standard protocol (38). All strains were grown on solid nematode growth media (NGM) and fed with *E. coli* OP50. Worms were maintained at 15°C unless specified. Hermaphrodite animals were used for the experimental purpose unless indicated otherwise. The specific life stage for each experiment is mentioned in the figure legend. Brood size, alae defect, gonadal arm migration and lifespan assays were done at 25°C after keeping the worms at 25°C for at least 3 generations. *alg-1(null)* and *alg-1(null); alg-2(CD)* animals were grown at 20°C for 2-3 generations and transferred at the L1 stage to 25°C for one generation to do phenotypic analysis.

### Mutant generation by CRISPR-Cas9 system

Genome editing of *C. elegans* by CRISPR-Cas9 method was done as previously described (39). F1 heterozygote and F2 homozygote mutants were identified using PCR genotyping and Sanger sequencing. All newly generated strains by CRISPR-Cas9 were outcrossed twice at least. All the strains used for this study and details of primers and oligonucleotide sequences for genome editing are listed in Table S1-S5.

### RNAi Knockdown

The knockdown of *alg-1* and *alg-2* in animals was carried out using the RNAi feeding method (37, 40). Synchronized L1 population (about 100 worms per NGM plate containing 100mM IPTG in triplicate) was exposed to bacteria expressing either control (L4440 plasmid), *alg-1* or *alg-2* specific dsRNA and maintained at 15°C and 25°C for about 50hrs and 117hrs, respectively.

### Lifespan assay

All strains were synchronized by treatment with an alkaline hypochlorite solution. For the lifespan assays, 25 to 30 L1 animals were plated on NGM plates containing OP50 and grown at 25°C. At the L4 stage, animals were transferred to new NGM plates with OP50 and FUDR (50µM). Adult worms were transferred every 2 days during active reproduction and scored for viability. Animals were scored as dead when they stopped responding to gentle prodding with a platinum wire pick and subsequently removed from the plate. Animals that died by internal hatching, vulval bursting, or crawling on the side of the plates were censored from the lifespan analysis. The Gehan-Breslow-Wilcoxon method was used to calculate *p-values* as there is a greater number of early death instance in mutant genotypes.

### Sperm quantification

Argonaute CD mutants were crossed with strain (RF1025), carrying a GFP-tagged histone transgene, *stls10027*, to quantify the spermatids. L4 hermaphrodites were upshifted from 20°C to 25°C, and their progeny was analyzed for the number of sperm they generated. Sperm quantification was completed following the outline mentioned in Huang et al., 2012 and Maniates et al., 2021 (41, 42), where L4m+4-hour hermaphrodite animals were smashed on a slide in sperm medium and then GFP positive spermatids were counted.

### Embryos and Young adults collection

All strains were synchronized by alkaline hypochlorite treatment and later suspended in M9 solution (22mM KH2PO4, 22mM Na2HPO4, 85mM NaCl, 1mM MgSO4). The embryos were collected for immunoprecipitation and maintained on NGM plates containing OP50 at 25°C for at least 3 generations before collecting young adults. The worm pellets were snap-frozen in Trizol for RNA extraction, while for Protein extraction, the worm pellets were snap-frozen in M9.

### Protein extraction

Extracts were prepared from synchronized young adult animals as well as embryos. Worms suspended and frozen in M9 solution were washed and resuspended in cold lysis buffer (10mM Potassium Acetate, 25mM HEPES-Potassium Hydroxide [pH 7.0], 2mM Magnesium Acetate, 1mM DTT, 0.5% [v/v] Triton X-100 and protease inhibitors) before being lysed by shaking with bead lysis kit (Next Advance RINO) using bullet blender Storm 24 (Next Advance). Then, the extracts were centrifuged at 17, 000g for 20 minutes at 4°C, and the clarified supernatants were collected and processed for Western Blot or Immunoprecipitation.

### Immunoprecipitation

Protein quantification was done using Bio-Rad Protein Assay (catalogue #500-0006). For immunoprecipitation, Dynabeads protein-G (Thermo Fisher Scientific, catalogue #10004D) was washed three times using cold lysis buffer and then incubated with rabbit polyclonal ALG-1 antibody in 250µl lysis buffer for 1 hr at room temperature with rotation. Antibodies bound to Dynabeads were washed thrice with lysis buffer and then incubated with protein extract overnight at 4°C with rotation. For endogenous flag-tagged ALG-2 immunoprecipitation, we used anti-FLAG M2 magnetic beads (Millipore Sigma, catalogue #M8823) directly.

### Western blotting

Equal amounts of proteins or beads were resuspended in 2X SDS loading buffer (100mM Tris-HCl [pH 6.8], 4% [w/v] SDS, 200mM DTT and 20% [v/v] glycerol) and then heated at 95°C for 10 min. Samples were resolved onto an 8% SDS-PAGE and transferred to Protran Premium NC membranes (GE Healthcare). The membranes were blocked with milk (5% [w/v] dried milk in 1X PBST (137mM NaCl, 2.7mM KCl, 10mM Na2HPO4, 1.8mM KH2PO4, 0.1%[w/v] Tween-20, [pH-7.2])) and later, incubated overnight at 4°C with primary antibodies. Rabbit polyclonal anti-ALG-1 and anti-AIN-1 antibodies were used at 1:1000 and 1:2000 dilution, respectively, in 1% [w/v] dried milk-supplemented PBST. Monoclonal mouse anti-beta-ACTIN (Abcam, catalogue #ab49900) antibody was used at 1:20000 dilution in 1% [w/v] dried milk supplemented PBST. Anti-FLAG (Millipore Sigma, catalogue #F1804) antibody was used at 1:10000 dilution in 1% [w/v] bovine serum albumin supplemented PBST. Membranes were then incubated with appropriate secondary antibodies before developing with Perkin-Elmer ECL Western Lightning Plus solution and visualized using a Chemidoc imaging system (Bio-Rad).

### Assessment of Argonaute expression and proteasomal degradation

Synchronized young adult worms were treated by rotating them in suspension for two hours with vehicle only (DMSO) or with 100µM MG132 (Sigma-Aldrich, catalogue #474790) in an M9 buffer. Then, worms were collected and washed in M9 before being lysed in the SDS loading buffer.

### Slicing assay

A specific target RNA probe (miR-1 target) was designed (Figure: S1A), and a 50 pm RNA probe was incubated with 20U T4 Polynucleotide kinase (PNK) (NEB, catalogue # M0201S), 1X T4 PNK buffer and 30 µCi P^32^ATP at 37°C for 30minutes (reaction volume 50µl) for radiolabelling reaction. Then, the reaction mix was incubated at 65°C for 20 minutes for heat inactivation. The reaction mix was purified using MicroSpin G-25 columns (Amersham, catalogue #27-5325-01).

Young adult worms lysed in cold lysis buffer (100mM Potassium Acetate, 30mM HEPES [pH 7.4], 2mM Magnesium Acetate, 1mM DTT, 0.5% [v/v] Triton X-100, protease inhibitors) were processed further for ALG-1 immunoprecipitation as detailed before. The IPed ALG-1 was then washed with slicing buffer (30mM HEPES [pH 7.4], 40mM Potassium Acetate, 5mM Magnesium Acetate, 5mM DTT, phosphatase inhibitor (Sigma, catalogue #4906845001)). Next, IPed ALG-1 was incubated in a slicing buffer containing a radiolabelled miR-1 target RNA probe along with Yeast RNA (Ambion, catalogue #AM7120G) and RNAse inhibitor at 20°C for 90 minutes for slicing reaction. When indicated, 1mM EDTA was added as an enzymatic inhibitor for the assay. Then, 2X PK buffer (0.2M Tris-HCl [pH 7.5], 25mM EDTA, 0.3M NaCl, 2% [w/v] SDS) was added to stop the reaction. Thereafter, RNA isolation was done using Trizol (Millipore Sigma, catalogue #T9424). Isolated RNA was run on UREA-PAGE, and the gel was exposed to a phosphorimager screen and visualized with Typhoon Imager (Amersham).

### RNA isolation and Northern Blot

Trizol (Millipore Sigma, catalogue: T9424) was used to extract total RNA from frozen worm pellets or beads after immunoprecipitation. Before Trizol extraction, those beads were suspended in 2X PK buffer (100 mM Tris-Cl [pH 7.5], 200 mM NaCl, 1% [w/v] SDS) and digested with proteinase K at 50°C for 20 minutes. 10-12 µg of total RNAs were used for northern blot. Samples were mixed with an equal volume of 2x Loading Dye (8M urea, 25 mM EDTA, 0.025% [w/v] xylene cyanol, 0.025% [w/v] bromophenol blue) and heated for 5 minutes at 80°C. 15% Urea Gel PAGE (Sequagel solutions) was pre-ran for 25 minutes before loading the samples. The gel was transferred onto a GeneScreen Plus Hybridization Transfer Membrane (Perkin Elmer, catalogue #NEF 988001PK). The RNA was cross-linked to the membrane with an EDC solution [0.373g (1-ethyl-3-(3- dimethylaminoprophy) carbodiimide and 1x methylimidazole (127.5mM 1- methylimidazole-HCl [pH 8]) at 60°C for 1 hour. The membrane was then washed with water several times and baked at 80°C for 10 minutes. The membrane was pre-hybridized in a hybridization bottle with 25 mL (5X SSC, 20mM Na2HPO4 [pH 7.2], 7% [w/v] SDS, 2X Denhardt’s Solution and 1mg of freshly denatured sheared salmon sperm DNA) at 50°C for 2.5 hours with rotation. The probes (found in Table S6), radiolabelled using Reaction Buffer (10mM Tris-HCl, 5mM MgCl2, 7.5mM DTT [pH 7.5]) and Exo-Klenow DNA polymerase (NEB, catalogue #M0212S); and Stop Solution (10mM EDTA) (IDT StarFire protocol) were heated at 85°C for 5 minutes and was directly added to the membrane and incubated in fresh pre-hybridization solution overnight at 50°C with rotation. The membrane was washed three times in a stringent wash solution (3X SSC, 5% [w/v] SDS) and once with a non-stringent wash solution (1X SSC, 1% [w/v] SDS) at 50°C with rotation. A phosphorimager screen was exposed to the membrane and revealed by autoradiography with Typhoon Imager (Amersham). Membranes were stripped by adding 100 mL of boiled 0.5% [w/v] SDS solution and incubating at room temperature twice for 10 minutes, and they were re-used again for hybridization after confirming the removal of the probe by exposure.

### miRNA cloning, sequencing, and analysis

We used the NEBNext multiplex small RNA library kit (catalogue #E7560S) for small RNA library preparation from young adults’ total RNA extracts and IPed samples (samples listed in Table S7-S8).

The small RNA sequencing data are available on Gene Expression Omnibus (GEO) with the accession number: GSE222733. Hiseq 4000 SR50 sequencing reads were mapped to the genome and cDNA using custom PERL (5.10.1) scripts and Bowtie 0.12.7 (43). Databases used include *C*. *elegans* genome (WormBase release WS215), Repbase 15.10 (44), and miRBase 16 (45). The Generic Genome Browser (46) was used to visualize the alignments. The samples were normalized to the total small RNAs (normalized reads can be found in Table S9-S10). All miRNA sequencing analysis was performed using R Statistical Software (v4.2.1; R Core Team 2022). Differential expression analysis was performed using Rankprod package (v3.22.0) and miRNAs with *p-value* ≤ 0.05 were considered significant. The log2 Normalised Reads Per Million are Z score transformed for generating Heatmaps with ComplexHeatmap package (v2.12.1). Spearman rank correlation was visualized using the Corrplot package (v0.92).

### miRNA detection by RT-qPCR

RT-qPCR of isolated total RNA was performed using TaqMan miRNA assay reagent (Life Technologies). ΔΔCT value was calculated using Sn2841 and U18 snoRNA as endogenous controls.

## RESULTS

### miRNA-specific slicing Argonautes are essential for animals’ fitness maintained in stressful conditions

To investigate the role of the endonuclease-slicing activity of miRNA-specific Argonautes in animals, we utilized CRISPR-Cas9 gene editing to replace the first aspartate with alanine in the catalytic tetrad on endogenous *alg-1* and *alg-2 loci (the other miRNA-specific Argonaute, ALG-5,* does not carry the essential residues needed for endonucleolytic cleavage and hence was not mutated (Figure 1A-B))*. As reported in several previous studies* (*27, 28, 30, 32, 47*), this point mutation is shown to abrogate the slicing activity of ALG-1 (Figure S1; As for ALG-2, despite multiple tests with different miRNA/RNA targets combination, we have not been able to see a robust cleavage activity from an immunoprecipitated complex, likely because the amount of miRNA loaded complex is clearly less than ALG-1). When we abolished the slicing activity of the Argonautes, we observed that this mutation did not affect the steady-state levels of ALG- 1 and ALG-2 (Figure S2). Besides animals expressing either ALG-1 or ALG-2 catalytic dead (CD) Argonautes, we also generated different double mutant animals to take into consideration the partial redundancy seen for *alg-1* and *alg-2* (33, 44–47).

We first monitored animals within the population for each genotype that displayed health issues (such as developmental arrest, egg-laying defect and protruding vulva) at the standard growing conditions (20°C). We observed that only animals expressing ALG- 2(CD) in *alg-1(null)* background were the only ones showing reduced fitness to a level greater than seen in *alg-1(null)* animals (Figure 1C, S3A). As an increased growing temperature creates physiological stress (48, 49), we next surveyed the phenotypes of the different mutant animals by maintaining them for at least three generations at 25°C.

In this specific condition, while single mutants, *alg-1* and *alg-2* catalytically dead animals displayed no significant health issues. However, animals expressing both ALG-1 and ALG-2 slicing dead Argonautes were found to be sick, with a portion of these mutants dying before reaching adulthood mainly due to larval death and bursting (Figure 1D, S3B). We also detected that the animals expressing ALG-1(CD) in the absence of wildtype *alg- 2* had significant defects, notably prominent egg-laying defects in the population (Figure 1D, S3B. As animals with *alg-1(null)* background cannot survive at 25°C, we were unable to score any mutant animals at this temperature (Not Determined (N.D.))). Thereafter, we pursued the phenotypic and molecular characterizations of ALG-1 and ALG-2 catalytically dead mutant animals, maintained at 25°C unless specified otherwise.

Next, we monitored the brood size of those mutants and observed a significant decrease for animals expressing either one or two miRNA-specific mutated Argonautes compared to wild-type (Figure 1E). The quantification of sperm and unfertilized oocytes of animals with significantly low brood size indicates that the decrease in progeny is not caused by a decline in sperm number or fertilization defects (Figure S4). Instead, it is more likely associated with the defect(s) occurring during oogenesis or embryogenesis. We conclude that the loss of the catalytic activity of both miRNA-specific ALG-1 and ALG-2 alters fitness and fecundity in *C. elegans* when maintained at more stressful growing conditions.

### The slicing activity of ALG-1 and ALG-2 pleiotropically influences miRNA function in animals

To specifically determine the role of the slicing activity of ALG-1 and ALG-2 in the miRNA function, we focus on phenotypes known to be affected by the alteration of this regulatory pathways in *C .elegans*. Among the developmental processes controlled by miRNAs, the alae is a cuticle structure, generated by the seam cells at the L4 to adult transition for which the proper timing and differentiation of these cells, during larval development, requires specific miRNAs, including *lin-4* and *let-7* miRNA family (34, 50–54). Any alteration of these miRNA functions would affect their number or cause defective fusion of these seam cells, leading to phenotypical incomplete alae breaks along the structure (55–57). Compared to wild-type animals, single and double mutants of catalytic dead Argonautes showed a significant increase in alae defects, and the effect of ALG-1(CD) is exacerbated by the loss of wild-type ALG-2 (Figure 2A). This data suggests that abrogating the enzymatic activity of miRNA-specific Argonautes affects the function of miRNAs in this developmental process.

**Figure 2.**
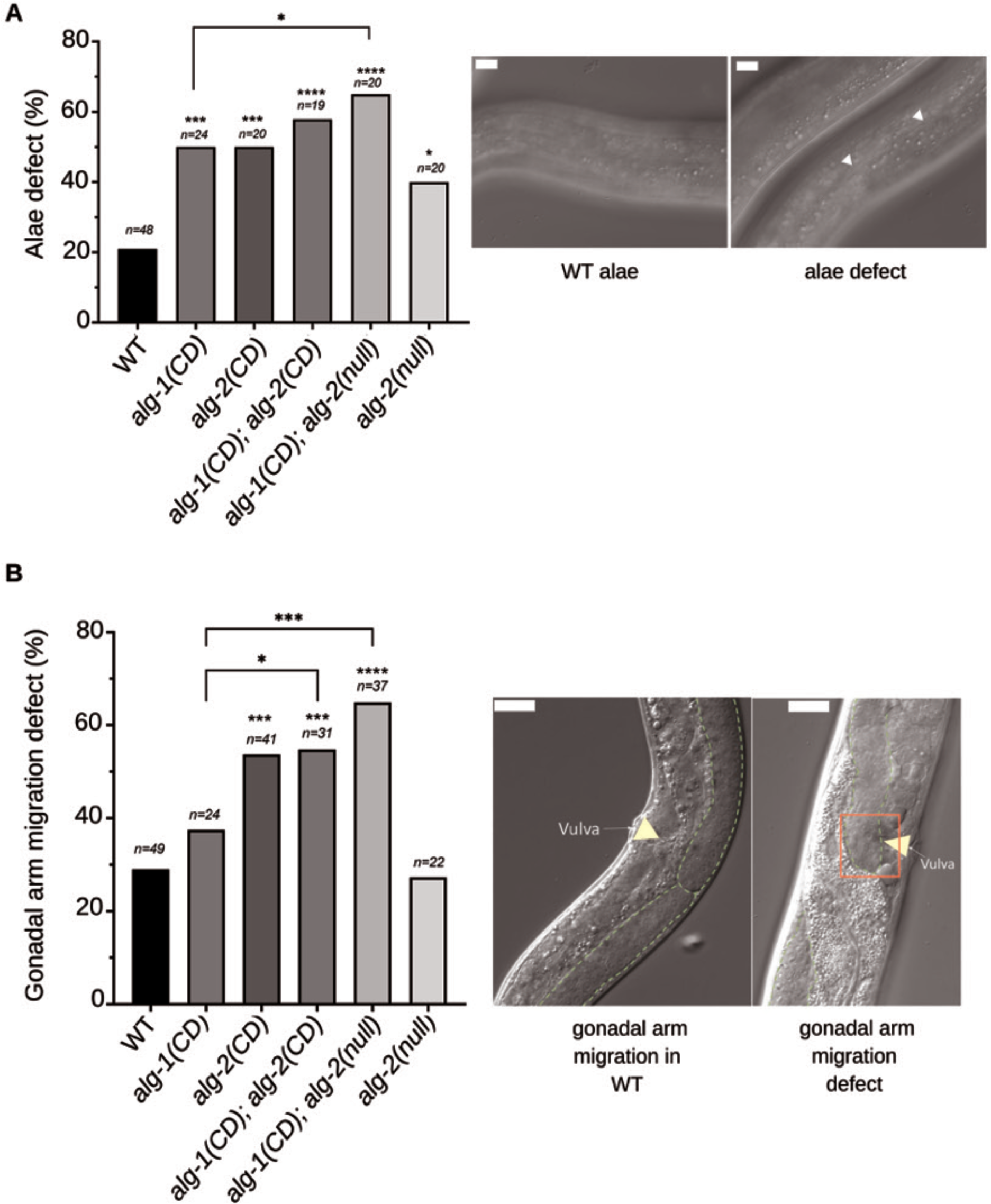
ALG-1 and ALG-2 catalytic activity contributed to the miRNA pathway. (A) Alae defects (indicated by arrows) and **(B)** gonadal arm migration defects in young adult animals, were scored under DIC Nomarski microscopy. The number of animals scored (*n*) is indicated. *p-value* was obtained by two tailed Fisher’s exact test (\**p*<0.05, \*\**p*<0.01, \*\*\**p*<0.001, \*\*\*\**p*<0.0001). Right side panels show representative images of defect observed on affected animals. The scale bars on alae and gonadal arm migration indicate 10µm and 20µm, respectively.

Another anatomical structure that requires miRNA activity is the hermaphrodite gonads of *C. elegans*. In this case, two miRNAs, miR-34 and miR-83, contribute to the appropriate migration of distal tip cells (DTC) of gonadal arms (58, 59) to create the distinctive U-shape structure found in adult animals. We observed that while the loss of ALG-1 catalytic activity alone does not influence the proper migration of animal gonads, the catalytic activity of ALG-2 is required for this process (Figure 2B). Since the loss of both ALG-1 and ALG-2 slicing activity does not cause an additive effect, we conclude that ALG-2 slicing activity is the most important contributor to this process.

It has been reported previously that ALG-1 (53, 60, 61) and specific miRNAs (62–68) are implicated in *C. elegans* lifespan regulation. We, therefore, wanted to determine whether the catalytic activity of ALG-1 and ALG-2 is involved in animal aging. As reported before (60), we observed that only the loss of *alg-1* but not *alg-2* leads to a shortened lifespan (Figure 3A-B). When we surveyed the single *alg-1* and *alg-2* catalytically dead mutants, we did not observe any significant effect on lifespan, but we noticed a significant reduction in lifespan in animals without any miRNA-specific slicer Argonautes (Figure 3C-G). Altogether, our data suggest that the slicing activity of ALG-1 and ALG-2 contributes to specific miRNA functions in animals.

**Figure 3.**
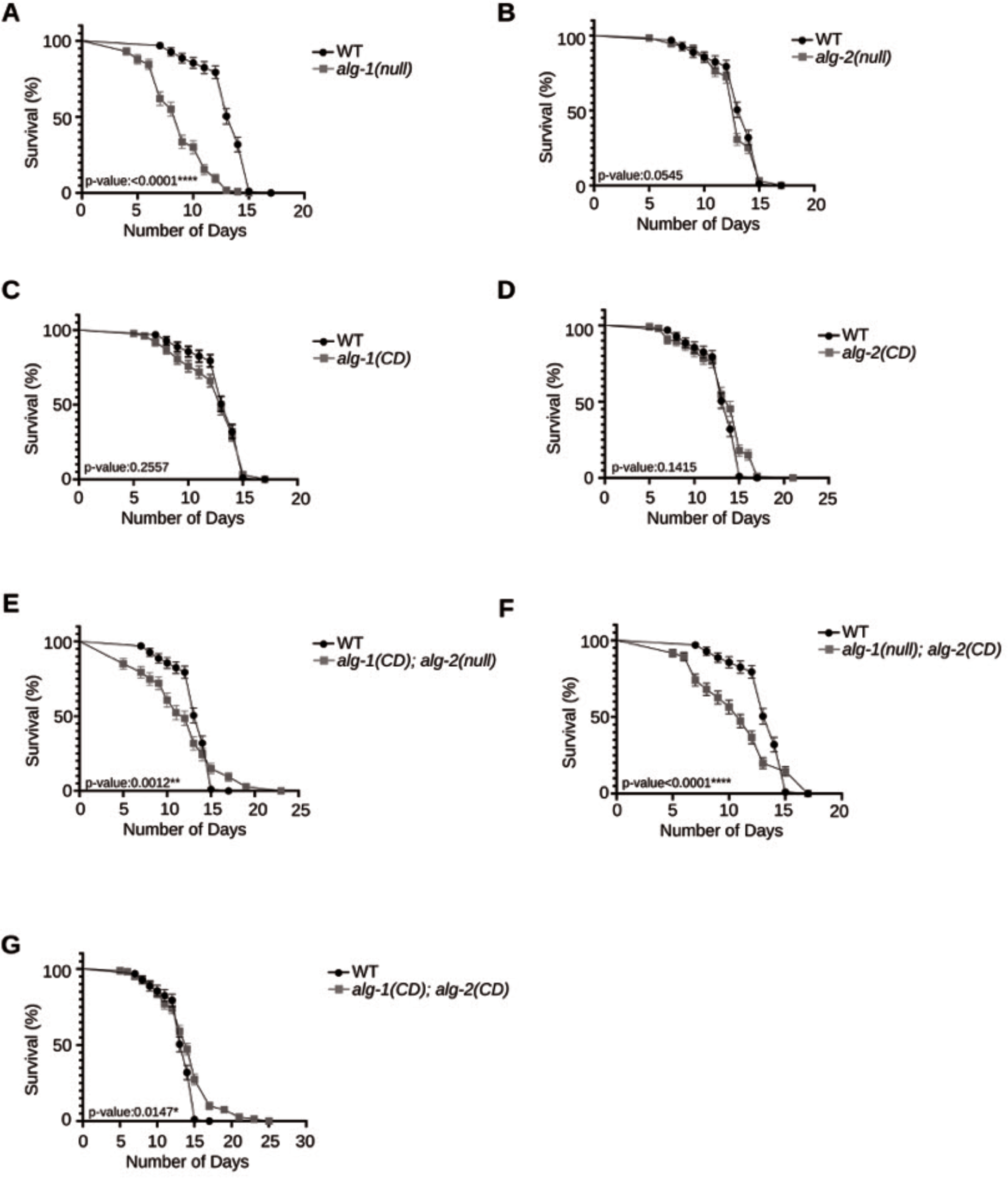
The role of slicer activity of ALG-1 and ALG-2 on animal lifespan. (A-G) Survival curve shows the lifespan of wild-type and ALG-1 and ALG-2 mutants. Lifespan assays were carried out with a population of *n*≥100 animals for each genotype. Same wild type data has been used to compare each mutant. *p-value* was obtained by Gehan- Breslow-Wilcoxon method (\**p*<0.05, \*\**p*<0.01, \*\*\**p*<0.001, \*\*\*\**p*<0.0001)

### Catalytic dead ALG-1 and ALG-2 trigger dysregulation of specific miRNAs in young adults

Previous studies in mice have shown that two erythroid-specific and atypical miRNAs necessitate the slicing activity of Ago2 to be correctly processed (27, 28, 30). To determine the broader contribution of miRNA-specific Argonaute in animals, we measured the abundance of miRNAs in young adults, carrying *ALG-1* and *ALG- 2* catalytically dead Argonautes by high-throughput sequencing.

A total of 176 miRNAs were detected, and those exhibiting near zero variance across all samples were excluded. Ultimately, 162 miRNAs were taken for further analysis (Figure 4, S5). Upon comparing the total miRNA profiles of each genotype, we found that *alg- 1(CD)* and *alg-1(CD); alg-2(CD)* animals exhibited similar expression profiles when compared to either *alg-2(CD) or wild-type* animals (Figure 4). Among the genotypes, the *alg-1(CD)* and *alg-1(CD); alg-2(CD)* animals were the only samples showing a significant positive Spearman correlation with each other (Figure S6A). The correlation and similarity in the expression pattern persisted even when the analysis focused on the significantly dysregulated miRNAs (Figure 5A; Figure S6B). This suggests that the loss of either ALG-1 or ALG-2 catalytic activity results in a marked difference in the expressed miRNA pool in animals. However, when both ALG-1 and ALG-2 activities are lost, there are more similarities in affected miRNAs with *alg-1(CD)* than *with alg-2(CD)* animals.

**Figure 4.**
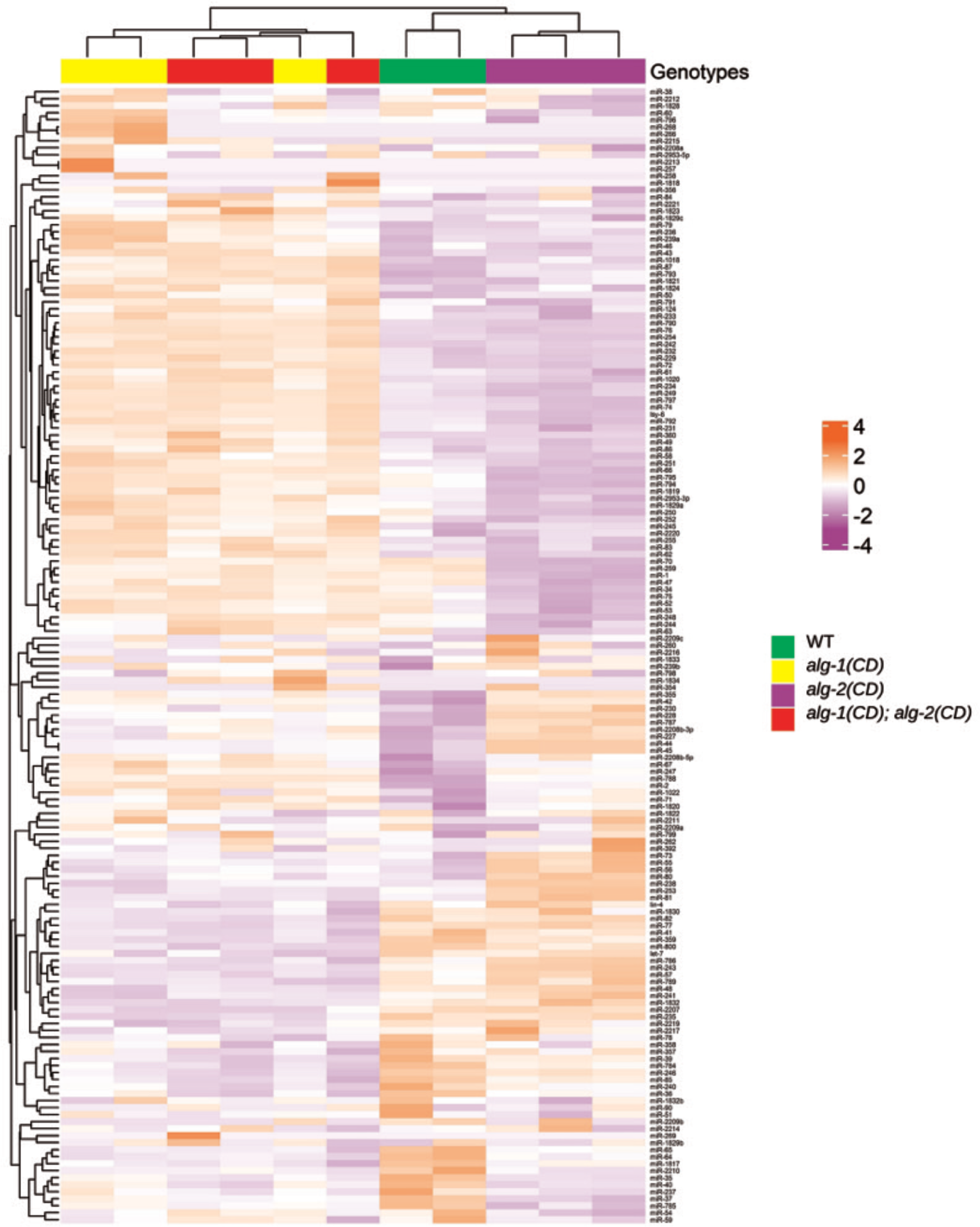
The loss of ALG-1 and ALG-2 slicing activity causes dysregulation in overall miRNA expression in young adults. Heatmap depicting the degree of similarity of each sample and their genotype based on the overall miRNA guide strand expression in young adults. *alg-1(CD)* and *alg-1(CD);alg-2(CD)* genotypes cluster together indicating their similarity towards each other and cluster away from *alg-2(CD)* and WT, indicating their dissimilarity towards the other two genotypes. The tile color represents the number of standard deviations above (Red) or below (Blue) the mean expression of a miRNA (Known as Z-Score). Agglomerative hierarchical clustering based on Euclidean distance is performed on the samples (columns) and miRNA (rows), which is represented by their respective dendrograms.

**Figure 5.**
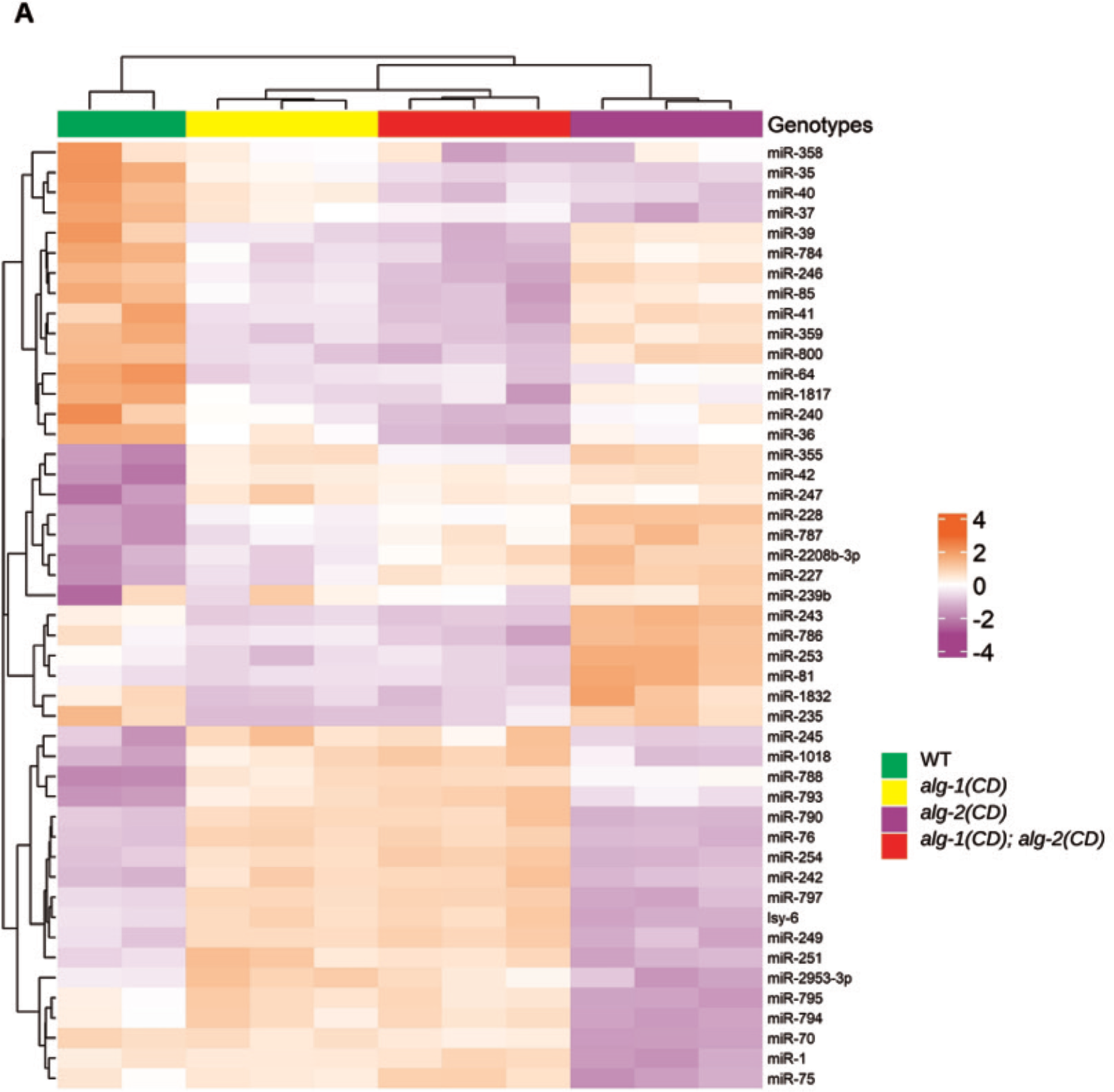
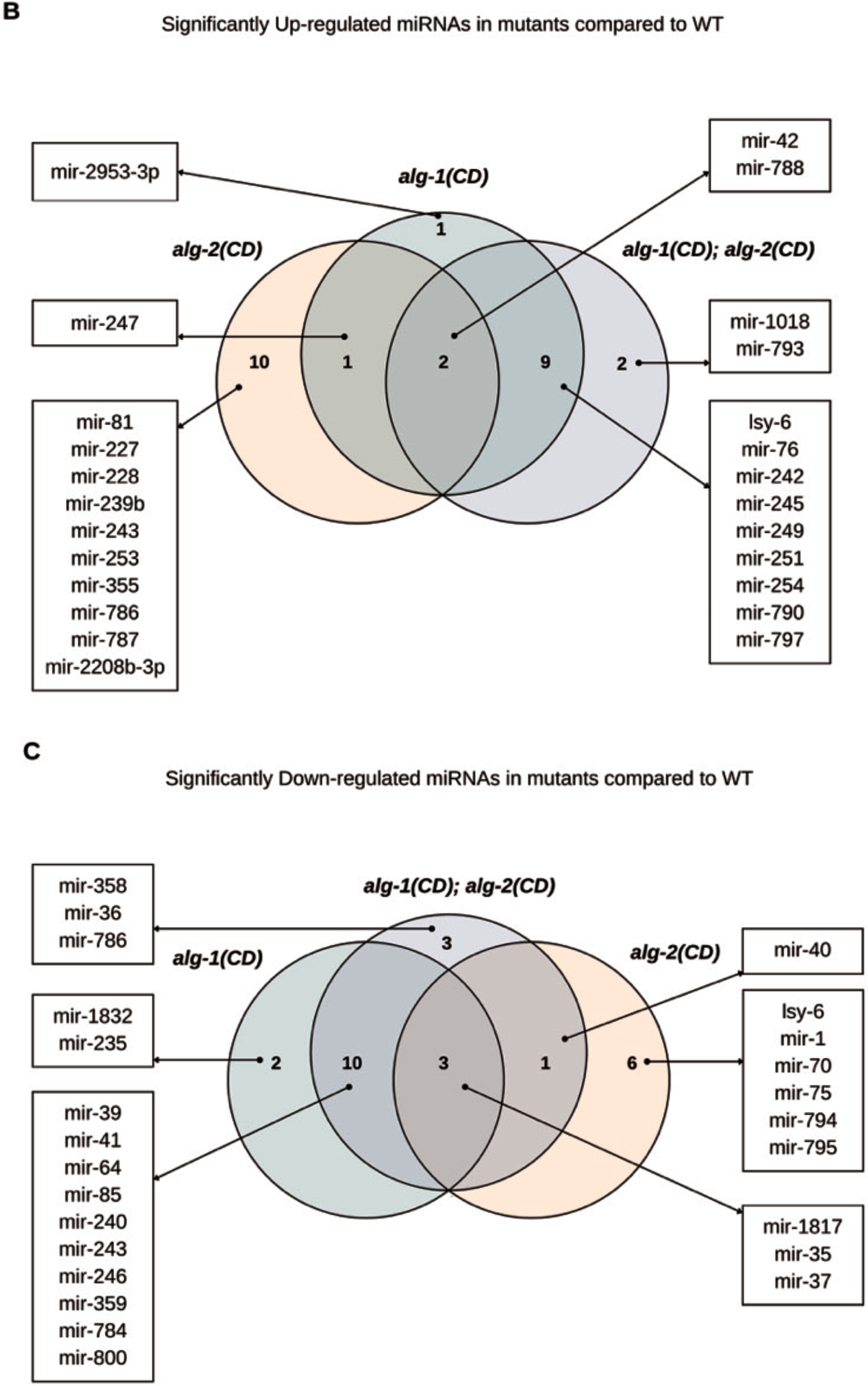
Catalytically dead ALG-1 and 2 Argonautes have significant and contrasting impact on the clusters of miRNAs. (A) Heatmap of the various clusters of significantly dysregulated miRNA (guide strand) in mutants compared to WT (2-3 replicates per strain). The tile color represents the number of standard deviations above (Red) or below (Blue) the mean expression of a miRNA (Known as Z-Score). Agglomerative hierarchical clustering based on Euclidean distance is performed on the samples (columns) and miRNA (rows), which is represented by their respective dendrograms. **(B)** Venn diagram depicting the significant up regulated miRNAs (guide strands) in *alg-1 (CD)* (light green), *alg-2(CD)* (light orange), *alg-1(CD); alg-2*(CD) (light blue) compared to WT. **(C)** Venn diagram depicting the significant down regulated miRNAs (guide strands) in *alg-1 (CD)* (light green), *alg-2(CD)* (light orange), *alg-1(CD); alg-2*(CD) (light blue) compared to WT.

When comparing all genotypes to the wild-type genotype, we identified 47 guide strand miRNAs that were significantly dysregulated. Specifically, in the single ALG-1 slicing dead mutant, we observed 28 significantly dysregulated miRNAs, with 24 of them also showing significant dysregulation in the double slicing dead mutant (Figure 5B-C). Conversely, in the single ALG-2 slicing dead mutant, we found 23 significantly dysregulated miRNAs, with only 6 displaying significant dysregulation in the double slicing dead mutant. (Figure 5B-C). To confirm the robust differences in the levels of the significantly dysregulated miRNAs, we also monitored some of them by TaqMan RT-qPCR (Figure S7). In contrast, the expression levels of star strand miRNAs did not show such a striking trend in dysregulation (Figure S8), which can be due to their low abundance, causing variability within replicates for each genotype. We, therefore, conclude that altering the slicing activity of ALG-1 and ALG-2 causes alteration in the level of specific subsets of miRNAs. A more significant overlap with affected miRNAs seen in the single *alg-1(CD)* animals suggests that the contribution of ALG-1 slicing activity on miRNA levels might be more preponderant than ALG-2 slicing activity in young adult animals.

To further gain some insights into the relationship between the miRNA changes and the observed phenotypes in the slicing dead mutants, we performed a Gene Set Enrichment Analysis (GSEA) and Over Representation Analysis (ORA) adapted for putative miRNA targets using miEAA 2.0 (69). While there was no enrichment in any known gene sets or pathways, we do note a significant over-representation of 4 putative target genes (*camt- 1*, *T22B11.4*, *AH9.6* and *F42H10.5*) for the significantly downregulated miRNAs in the *alg-1(CD); alg-2(CD)* double mutant only (Figure 5C; Table S11). This could suggest that these target genes are upregulated within the slicing dead mutants. However, the same genes are found in the single ALG-1 and ALG-2 mutants but are not significantly overrepresented, suggesting that these genes could be dysregulated in all the slicing dead mutants but to varied degrees.

Next, to determine if miRNA production is affected by the absence of catalytic miRNA-specific Argonautes, we monitored the level of precursor and mature miRNA molecules by Northern blots. When we probed for specific miRNAs in animals expressing ALG-1 and ALG-2 CD Argonautes, we did not observe any accumulation of pre-miRNA precursors in any condition except for miR-37 (Figure S9). Thus, even though altering both ALG-1 and ALG-2 slicing activity affects the level of a subset of miRNAs, it does not cause either accumulation of miRNA precursor molecules (except for miR-37) as observed in the complete absence of Argonautes (57, 68) (Figure S9) or create truncated miRNA precursors as seen before when overexpressing catalytically inactive ALG-1 or ALG-2 (37, 54). It is important to note here that the mature miRNA, quantified during northern blot, does not necessarily follow the same pattern as observed during small RNA sequencing analysis because this experimental approach cannot distinguish miRNAs that share common sequences (like members of the same miRNA family) as the stringency for the probing is not so high for a 21nt long RNA, whereas sequencing can distinguish them. Altogether, the results suggest that the loss of ALG-1 and ALG-2 slicing activity has a contrasting effect on miRNA levels in animals that is not a consequence of a defect in their production.

## DISCUSSION

In this study, we explored the actual global contribution of slicer Argonautes in the miRNA pathway. Previous studies in mammals only showed that the slicer Argonaute Ago2 participated in the biogenesis of two distinct and atypical miRNAs, miR-451 and miR-486 (27–30). Importantly, they could not exclude the contribution of Ago3, another Argonaute that binds miRNAs with a slicing capacity (33). Furthermore, Ago2 is also needed in the mammalian system for other small RNA pathways, such as siRNAs (31, 32), which complicated the phenotypical analyses of the Ago2CD mutant. Thus, the studies so far have prevented the precise analysis of the contribution of slicing Argonautes to the miRNA pathway in animals. The utilization of *C. elegans* represents the ideal system for elucidating the function of slicing Argonautes in the miRNA pathway, as ALG-1 and ALG- 2, essential for miRNA-mediated gene regulation, are not involved in other small RNA pathways in this animal model (34, 35).

Our previous study has reported that the slicing activity of ALG-1 and ALG-2 was essential for animal viability (37). In contrast, our new data here supported the fact that the slicing activity of ALG-1 and ALG-2 has a more modest contribution in animals grown in optimized conditions. Furthermore, when we altered the expression level of wild-type ALG-1 and ALG-2 by RNAi into *alg-2(CD)* and *alg-1(CD)* mutant animals, respectively, in a similar manner as performed in the previous report (37), we did not phenocopy the strains overexpressing slicing dead ALG-1 and ALG-2 (Figure S10). The discrepancy between those two studies can likely be explained by the expression level of ALG-1CD and ALG-2CD proteins achieved with both experimental approaches. Bouasker and Simard’s analysis focused on the expression ALG-1 and ALG-2 slicing dead proteins using extrachromosomal arrays, which leads to a substantial overexpression of the protein. Here, we modified the endogenous loci using the CRISPR-Cas9 gene editing method, which retains the same expression level of endogenous proteins. Thus, the first study with overexpressing conditions likely affects slicing dead ALG-1 and ALG-2 interaction with miRNAs and protein partners, which could cause an imbalance in molecular stoichiometry, leading to a potential dominant effect. The mutations of the endogenous loci of *alg-1* and *alg-2* genes performed here prevent causing this effect.

The phenotypic analysis of *alg-1* and *alg-2* catalytically dead mutants demonstrate their contribution to different biological pathways regulated by miRNAs, such as fertility, embryogenesis, developmental timing, alae formation, and gonadal arm migration, that are noticeable when animals are maintained at a higher temperature. Interestingly, we observed that the effect on mutant animals with different genotypes varied, indicating a distinctive contribution of slicing ALG-1 and ALG-2 towards these processes and the restricted slicer function of ALG-1 or ALG-2 to specific cells, tissues, or developmental stages may consequently shape the impact on particular miRNAs. For instance, altering the ALG-2 slicing activity leads to a more penetrant effect on brood size than affecting the enzymatic function of ALG-1. We believe the more robust phenotype observed in *alg- 2(CD)* reflects its capacity to interact with miRNAs and proteins, thus likely creating non-functional complexes whereas in *alg-2* null animals, miRNAs and proteins that usually bind ALG-2 are now devoid of binding ALG-1, which can compensate for the loss of ALG-2. *On the other hand, the observed phenotypes are not as penetrant as the ones observed with a complete loss of either specific miRNA such as let-7 or lin-4 or both* alg- 1 *and* alg-2 *genes* (34, 50, 55, 56, 70, 71), it suggests that the slicing Argonautes are not essential for canonical miRNA function and the viability of the worms, at least in optimal growing conditions. Molecularly, our data indicated that while the slicing activity of ALG- 1 and ALG-2 does not affect its association with the GW182 protein, AIN-1 or miRNAs (Figure S11), it does have a significant effect on the level of specific subsets of miRNAs. Among the ones affected, the members of miRNA family(s) are not uniformly dysregulated in CD mutants, as seen for the miR-2, miR-35 and miR-251 families. This shows that even though the expression level of some miRNAs is significantly affected, other miRNA family members most probably contribute to the successful survival of CD mutant animals. Moreover, previous studies indicated a relationship between heat stress response and specific miRNAs (miR-64-66 cluster, miR-71, miR-80, miR-85 or miR-229) in worms (63, 67, 72, 73). It has been found that heat sensitivity, caused by the loss of the miR-64 family (miR-64-66 clusters, miR-229), influences worm survival negatively (72). Incidentally, we can see from our sequencing data that miR-64 is significantly downregulated in CD mutants compared to wild-type animals. This ascertains the notion that the slicing activity of ALG-1/2 dependent optimal expression of specific miRNA(s) is relevant for the survival of the animals at higher temperatures.

Among the significantly affected miRNAs, we cannot find one whose misregulation can explain the pleiotropic phenotypes observed in *alg-1(CD)* and *alg-2*(*CD)* animals. Furthermore, no significant gene sets or pathways are represented in the gene set enrichment analysis, suggesting dysregulation in multiple pathways or gene sets. Additionally, the 4 significantly overrepresented target genes identified from the significantly downregulated miRNA in the double mutant were, however, observed to be overrepresented in both the single ALG-1 and ALG-2 mutants. While the over representation analysis results from the individual ALG-1 and ALG-2 mutants are statistically nonsignificant, their presence in the analysis suggests that these 4 genes are not unique to a specific slicing dead genotype. In addition, we do also observe noticeable but not statistically significant changes in the levels of several miRNAs (such as lin-4 and let-7 family members) for which cumulative effect could cause phenotypes seen in mutant animals. We can conclude that the slicer activity contributes to maintaining the expression level of different miRNAs, and the consequence of altering their levels affects different cellular pathways whose combined effect leads to the observable phenotypes in animals exposed to more stressful conditions.

Interestingly, the McJunkin’s group have created independently the same slicing dead *alg-1* and *alg-2* alleles (see companion study). When we compared our miRNA sequencing data with the ones from Kotagama *et al.*, we did not observe the same molecular effects on miRNAs abundance. There are several points of difference in the worm maintenance as well as small RNA sequencing sample preparation and analysis that can explain the data discrepancy. Firstly, in our case, the animals are maintained at 15°C and shifted to 25°C for at least 3 generations before harvesting samples or phenotyping, while the Kotagama et al. study shifts the animals to 25°C from the 20°C maintenance temperature after synchronization (early L1). Secondly, our small RNA libraries are size-selected after PCR amplification, and the Kotagama et al. study performed a size selection immediately after reverse transcription prior to the PCR amplification of the libraries. Finally, the raw data following sequencing is analyzed by different pipelines. In our case, the Reads Per Million value of each sample was normalized to the total small RNAs sequenced, and then differential expression analysis was carried out using the Rank Product Method. In the companion study, the aligned counts of each sample are analysed using DESeq2 methodology. To rule out that the difference was technical and related to the pipeline analyses, we re-analyzed both datasets using the same pipelines (e.g. DeSeq2 and our described pipeline). Therefore, when analysis pipelines are the same, we see overlapping changes in miRNA abundances, and we conclude that the remaining differences seen between our respective studies are attributable to different growing conditions.

*In the past, in vitro* and *in cellulo* approaches have demonstrated that Argonaute slicing activity can help remove the star (or passenger) strand from the miRNA duplex created by Dicer cleavage (27, 47). Consistent with this model, Kotagama *et al*. observed a significant increase of a subset of star miRNAs in *alg-2(CD)* embryo samples. In contrast here, when we monitored the level of guide and star miRNA strand pairs of the significantly dysregulated miRNAs in young adults, despite their low abundance, we observed subsets of miRNA duplexes that appear to accumulate distinctively in *alg- 1(CD)* and *alg-2(CD)* animals (Figure S12). However, the variability detected for the miRNA star counts in wild-type samples (likely due to their low abundance) prevents us from assessing any significant effect of Argonaute catalytic activity on the miRNA duplex. Nevertheless, our data could suggest that in young adult animals, both ALG-1 and ALG- 2 slicing activity might contribute to duplex cleavage, while in embryos, ALG-2 appears to be the main contributor to this process.

Previous studies in culture cells and *C. elegans* using ectopically overexpressed slicing dead Argonautes have reported the production of atypical miRNA precursor molecules in the absence of catalytically active Argonautes (37, 47). With the approach used here that maintains the endogenous level of ALG-1 and ALG-2, we did not observe any aberrant form of miRNA molecules. This suggests that over-expressing slicing dead Argonautes can affect the miRNA processing machinery, leading to the production of aberrant molecules. Our data stresses the necessity of performing structure-function studies in experimental conditions in which the molecular stoichiometry of the proteins must be maintained. The recent development of powerful genome editing approaches allows us to design studies to uncover the precise gene function in critical gene regulatory pathways.

In summary, our study highlights that even though the slicing function is preserved in miRNA-specific Argonautes, their physiological and molecular contributions to the canonical miRNA pathway are discernible only in animals subjected to a more challenging environment, such as exposure to elevated temperatures. This is further supported when we compare our analysis with Kotagama *et al.* who conducted their experiment in less stressful conditions and did not observe any significant phenotypes in animals without miRNA-specific slicing Argonautes. Hence, we can visualize that the significance of slicing Argonautes becomes paramount in diverse conditions encountered by animals in the wild, where components or specific molecular steps involved in releasing the miRNA strand from duplexes may become less efficient. Future investigations testing this hypothesis will likely uncover the contributions of diverse underappreciated protein functions, such as slicing Argonautes, that cannot be fully elucidated yet under our well-controlled laboratory conditions.

## DATA AVAILABILITY

Raw and processed small RNA-seq datasets have been deposited at NCBI’s Gene Expression Omnibus (GEO) repository and are publicly available as of the date of publication. Accession number for the data is GSE222733. This paper does not report original code. All unique and stable reagents and strains generated in this study are available from the lead contact without restriction.

## FUNDINGS

This work was supported by The Canadian Institutes of Health Research (CIHR) (M.J.S.) and National Institutes of Health (NIH, GM126458) (A.L.A).

## ACKNOWLEDGMENTS

We would like to thank Dr. Katherine McJunkin for sharing data with us prior to publication. We also thank Dr. Weifeng Gu for the help with small RNA analyses, Sarah Côté for technical help, Dr. Zhirong Bao for sharing the *stls10027* transgene strain used for spermatid quantification and Dr. Andrew Singson for providing laboratory resources to complete the sperm quantification. Some strains used in this study are provided by *Caenorhaditis* Genetic Center, which is funded by NIH Office of Research Infrastructure Programs (P40 OD010440). We thank the members of our laboratory for helpful comments. The graphical abstract and the schematic diagram in Figure 1B were created with BioRender.com. A.P. received a scholarship from Université Laval. V.V. received a scholarship from the Faculty of Medicine of Université Laval and from the Fondation du CHU de Québec. M.L. was a recipient of a scholarship from Natural Sciences and Engineering Research Council of Canada (NSERC). K.A.M. received a NIH Institutional Research and Academic Career Development Award (K12GM093854) Fellowship. M.Q.H. was recipient of scholarship from the Fonds de Recherche du Québec-Santé (FRQ-S).

## AUTHOR CONTRIBUTIONS

Conceptualization, A.P. and M.J.S.; Methodology, A.P., V.V., M.L., F.H., K.A.M. M.Q.H., A.L.A. and M.J.S.; Investigation, A.P., V.V., M.L., F.H., K.A.M. and M.Q.H.; Writing –Original Draft, A.P. and M.J.S.; Writing – Review & Editing, A.P., V.V., M.L., F.H., K.A.M. M.Q.H., A.L.A. and M.J.S. Funding Acquisition, M.J.S.; Supervision, A.L.A. and M.J.S.

## DECLARATION OF INTEREST

The authors declare no competing interests.

## Notes

### Competing Interest Statement

The authors have declared no competing interest.

### Summary of Updates

- New figure panels 1C-E and 3A-G -New supplemental Figures S1, S2, S3, S5, S7, S9, S10 -Multiple text sections clarified

