## Supplemental Figures for "Defining the contribution of microRNA-specific slicing Argonautes in animals"

#### SUPPLEMENTAL FIGURES AND LEGEND

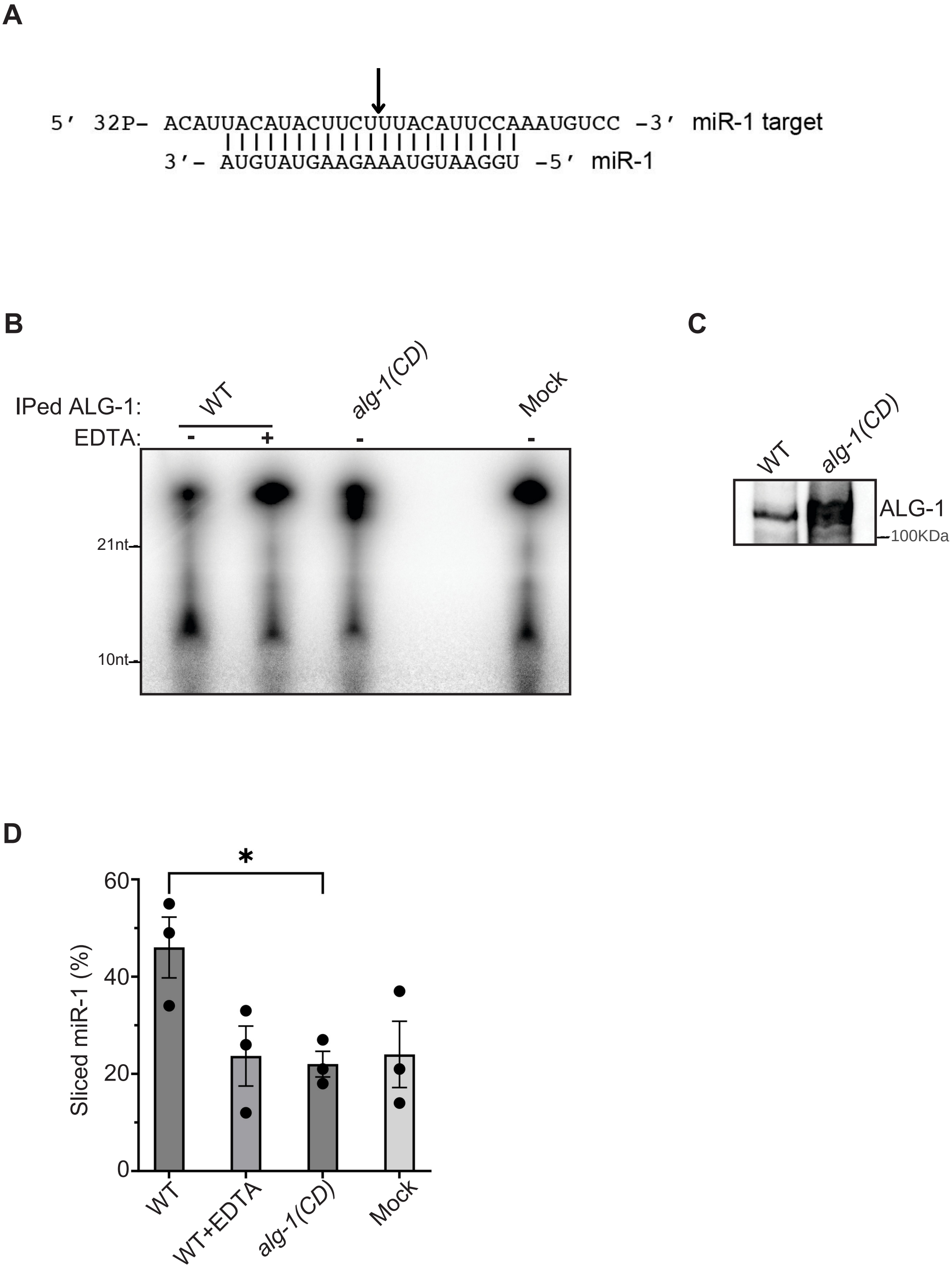
A
5′ 32P- ACAUUACAUACUUCUUUACAUUCCAAAUGUCC -3′ miR-1 target
3′- AUGUAUGAAGAAAUGUAAGGU -5′ miR-1
B
IPed ALG-1:
WT
alg-1(CD)
Mock
EDTA: - + - -
21nt
10nt
C
WT
alg-1(CD)
ALG-1
100KDa
D
*
60
40
20
0
Sliced miR-1 (%)
WT
WT+EDTA
alg-1(CD)
Mock

**Figure S1**

**Figure S1.** Mutation in catalytic tetrad of endogenous ALG-1 causes loss of its slicer function. **(A)** Diagrammatic representation of $P^{32}$ radiolabelled miR-1 target probe, used for slicer assay. **(B)** Representative UREA-PAGE blot (N=3) shows cleavage of radio-labelled miR-1 target probe in presence of immuno-precipitated wt and catalytically dead ALG-1, collected from young adult worm extract. EDTA has been used as an inhibitor of slicer assay. **(C)** Western blot shows the presence of ALG-1 in the IP-ed fraction used in **(B)**. **(D)** Quantification of cleaved radio-labelled miR-1 target probe (Cleaved/Total). The *P-value* (*$p<0.05$) was obtained by two tailed unpaired t-test.

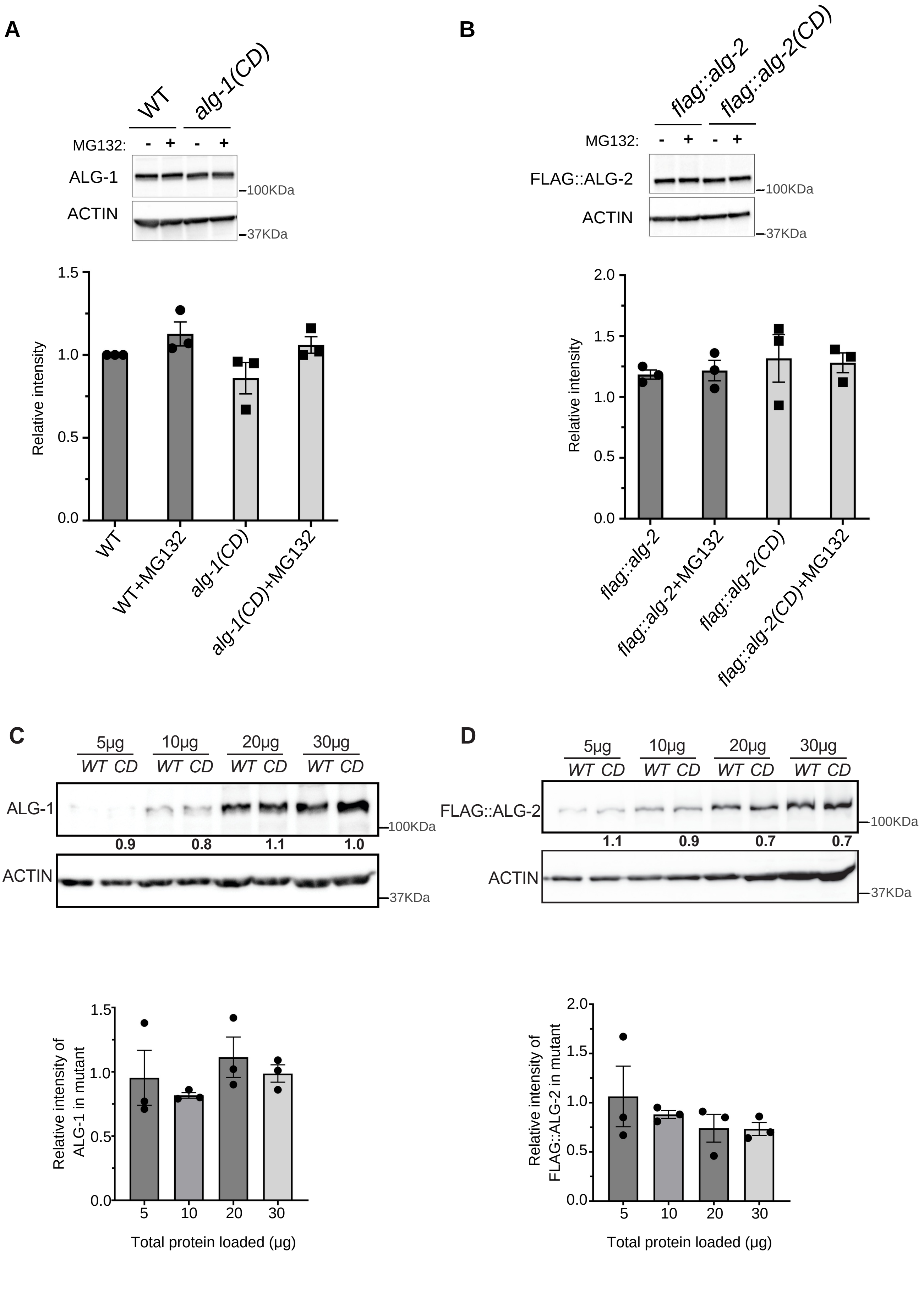
A
WT
alg-1(CD)
MG132: - + - +
ALG-1
100KDa
ACTIN
37KDa
Relative intensity
0.0
0.5
1.0
1.5
WT
WT+MG132
alg-1(CD)
alg-1(CD)+MG132
B
flag::alg-2
flag::alg-2(CD)
MG132: - + - +
FLAG::ALG-2
100KDa
ACTIN
37KDa
Relative intensity
0.0
0.5
1.0
1.5
2.0
flag::alg-2
flag::alg-2+MG132
flag::alg-2(CD)
flag::alg-2(CD)+MG132
C
5µg 10µg 20µg 30µg
WT CD WT CD WT CD WT CD
ALG-1
100KDa
0.9 0.8 1.1 1.0
ACTIN
37KDa
Relative intensity of ALG-1 in mutant
0.0
0.5
1.0
1.5
5 10 20 30
Total protein loaded (µg)
D
5µg 10µg 20µg 30µg
WT CD WT CD WT CD WT CD
FLAG::ALG-2
100KDa
1.1 0.9 0.7 0.7
ACTIN
37KDa
Relative intensity of FLAG::ALG-2 in mutant
0.0
0.5
1.0
1.5
2.0
5 10 20 30
Total protein loaded (µg)

**Figure S2**

**Figure S2. Steady state level and stability of ALG-1 and ALG-2. (A,B)** Western blots show the stability of ALG-1(CD) and endogenously FLAG tagged ALG-2(CD) compared to ALG-1 and FLAG::ALG-2 in presence of 100µM proteasome inhibitor (MG132) in young adults. The images represent one of the blots of the replicates (N=3). Relative intensity has been quantified from experimental replicates. **(C,D)** Western blots show dilution series for both ALG-1(CD) and endogenously FLAG tagged ALG-2(CD) compared to ALG-1 and FLAG::ALG-2. The images represent one of the blots of the replicates (N=3). Relative intensity (CD/WT) has been quantified from experimental replicates. The *P*-value was obtained by t-test.

**A**

Phenotype breakdown

| Temp. | Genotype | Total (*n*) | Dead (n) | EGL (*n*) | Sick and pale OR P.V. OR retarded develop-ment (*n*) | Molting (*n*) |
|---|---|---|---|---|---|---|
| 20°C | *wt* | 313 | 2 | - | 2 | - |
| | *alg-1(CD)* | 395 | 2 | - | - | - |
| | *alg-2(CD)* | 297 | - | 2 | - | - |
| | *alg-1(CD); alg-2(CD)* | 434 | 5 | 16 | - | 1 |
| | *alg-1(CD); alg-2(null)* | 291 | - | - | 1 | - |
| | *alg-1(null); alg-2(CD)* | 116 | 10 | - | 99 | - |
| | *alg-1(null)* | 238 | 14 | - | 103 | - |
| | *alg-2(null)* | 325 | - | - | 16 | - |

P.V.- Potruding Vulva

**Figure S3**

**B**

Phenotype breakdown

| Temp. | Genotype | Total (*n*) | Dead (*n*) | EGL (*n*) | Sick and pale OR P.V. OR retarded develop-ment (*n*) | Not wild type looking (*n*) |
|---|---|---|---|---|---|---|
| 25°C | *wt* | 582 | 2 | 15 | - | - |
| | *alg-1(CD)* | 303 | - | 2 | 1 | 10 |
| | *alg-2(CD)* | 182 | - | 26 | - | - |
| | *alg-1(CD); alg-2(CD)* | 178 | 11 | - | - | 167 |
| | *alg-1(CD); alg-2(null)* | 192 | - | 56 | - | - |
| | *alg-1(null); alg-2(CD)* | N.D. | - | - | - | - |
| | *alg-1(null)* | N.D. | - | - | - | - |
| | *alg-2(null)* | 512 | - | 26 | - | - |

P.V.- Potruding Vulva

#### Figure S3

**Figure S3. Phenotypic breakdown. (A,B)** The detailed breakdown of phenotypes observed in worms , grown at 20°C and 25°C.

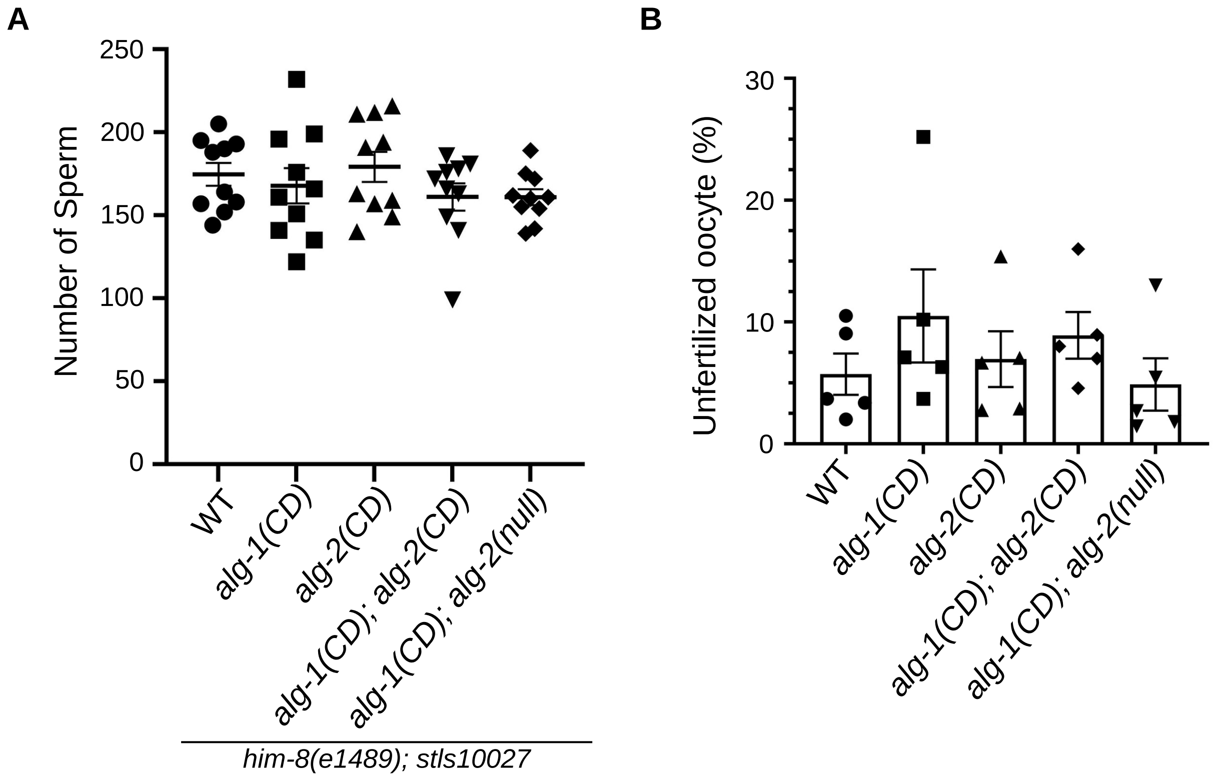


**Figure S4**

**Figure S4. The effect of ALG-1 and ALG-2 catalytic activity on fertility. (A)** Quantification of sperm, present in catalytic argonaute mutants in specific background. Mean along with SEM for each genotype are shown. The sperm are counted from *n*=10 worms for each genotype. One-way anova followed by Tukey's multiple comparison test was done to find out the significance between multiple genotypes. **(B)** Percentage of unfertilised oocyte was counted after putting 4 young adults per plate (*n*=5) and monitored for 48 hrs. The *P*-value was obtained by two tailed t-test.

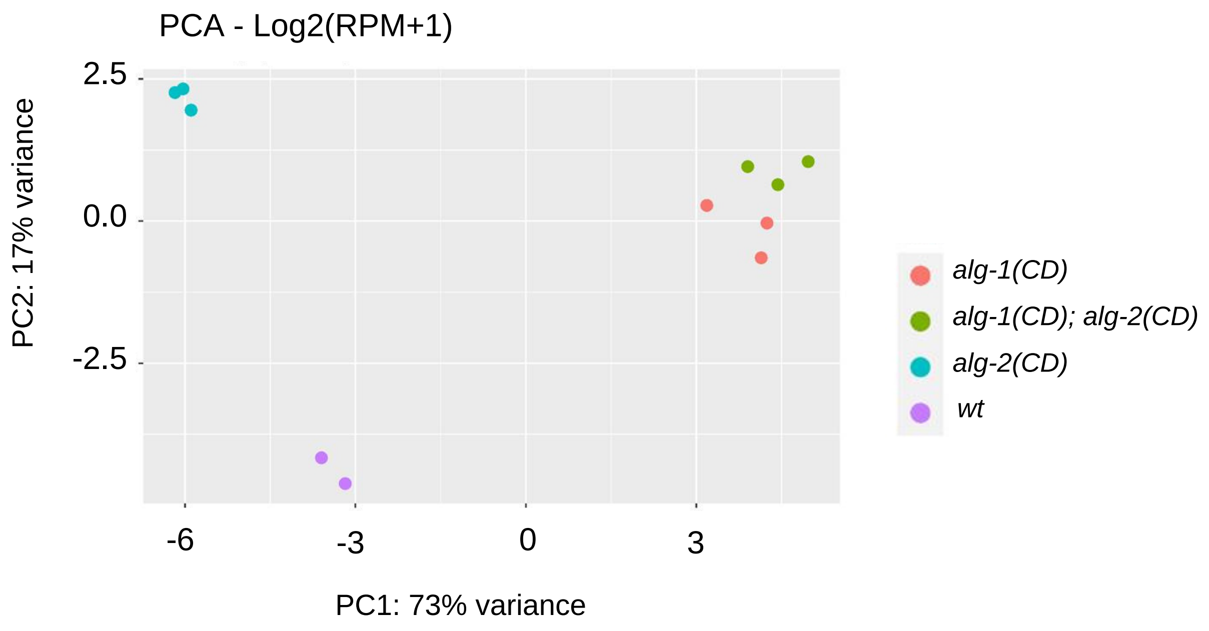


#### Figure S5

**Figure S5. PCA analysis.** Principal component analysis of the small RNA sequencing data using log2 Normalised Reads Per Million data of 162 micro-RNAs. The different genotype groups are represented by different colors as indicated by the legend provided within the graph and each dot of a genotype represents a replicate of a small RNA-seq sample.

A

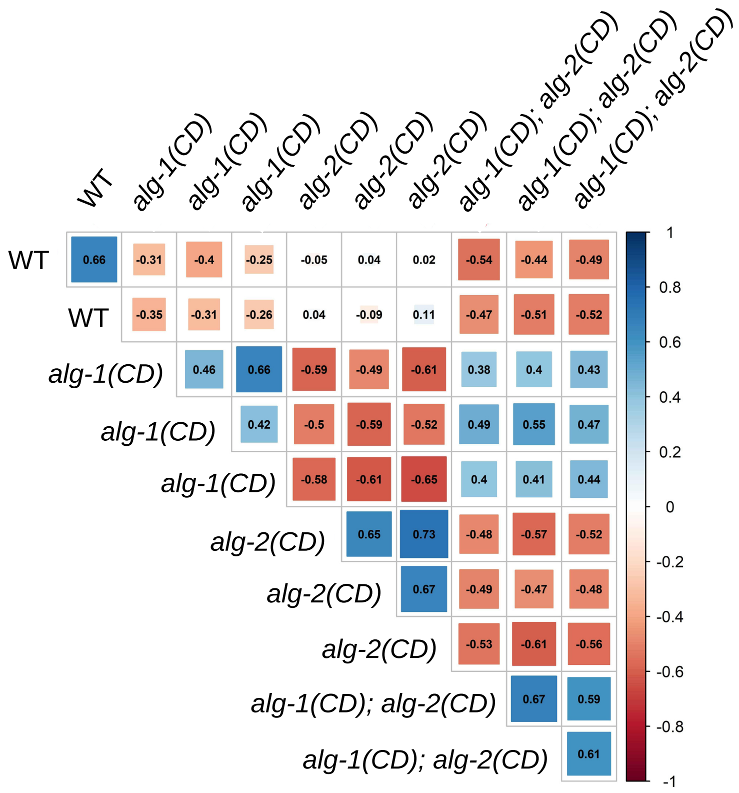
WT
alg-1(CD)
alg-1(CD)
alg-1(CD)
alg-2(CD)
alg-2(CD)
alg-2(CD)
alg-1(CD); alg-2(CD)
alg-1(CD); alg-2(CD)
alg-1(CD); alg-2(CD)

B

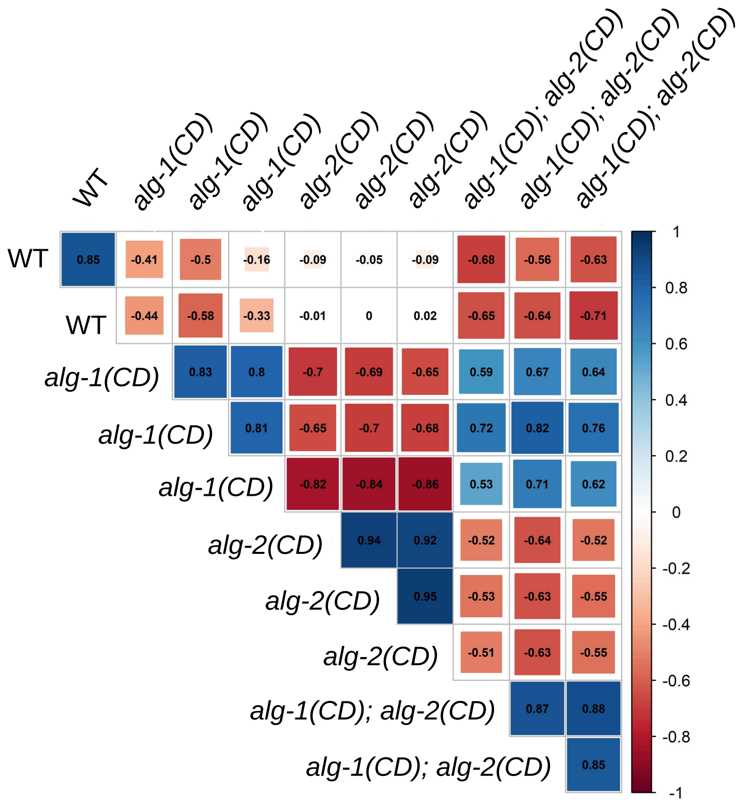
WT
alg-1(CD)
alg-1(CD)
alg-1(CD)
alg-2(CD)
alg-2(CD)
alg-2(CD)
alg-1(CD); alg-2(CD)
alg-1(CD); alg-2(CD)
alg-1(CD); alg-2(CD)

**Figure S6**

**Figure S6. Correlation among the samples and genotypes based on the mature miRNA population. (A)** Correlation performed using all 162 expressed miRNA guide strands with the spearman correlation coefficient (ρ) reported for each sample correlation with another. *alg-1(CD)* positively correlates to *alg-1(CD);alg-2(CD)* with a correlation value ranging from 0.38 to 0.55 **(B)** Correlation performed using 47 significantly dysregulated miRNA guide strands with the spearman correlation coefficient (ρ) reported for each samples correlation with another. *alg-1(CD)* shows increased positive correlation to *alg-1(CD);alg-2(CD)* with value ranging from 0.53 to 0.82.

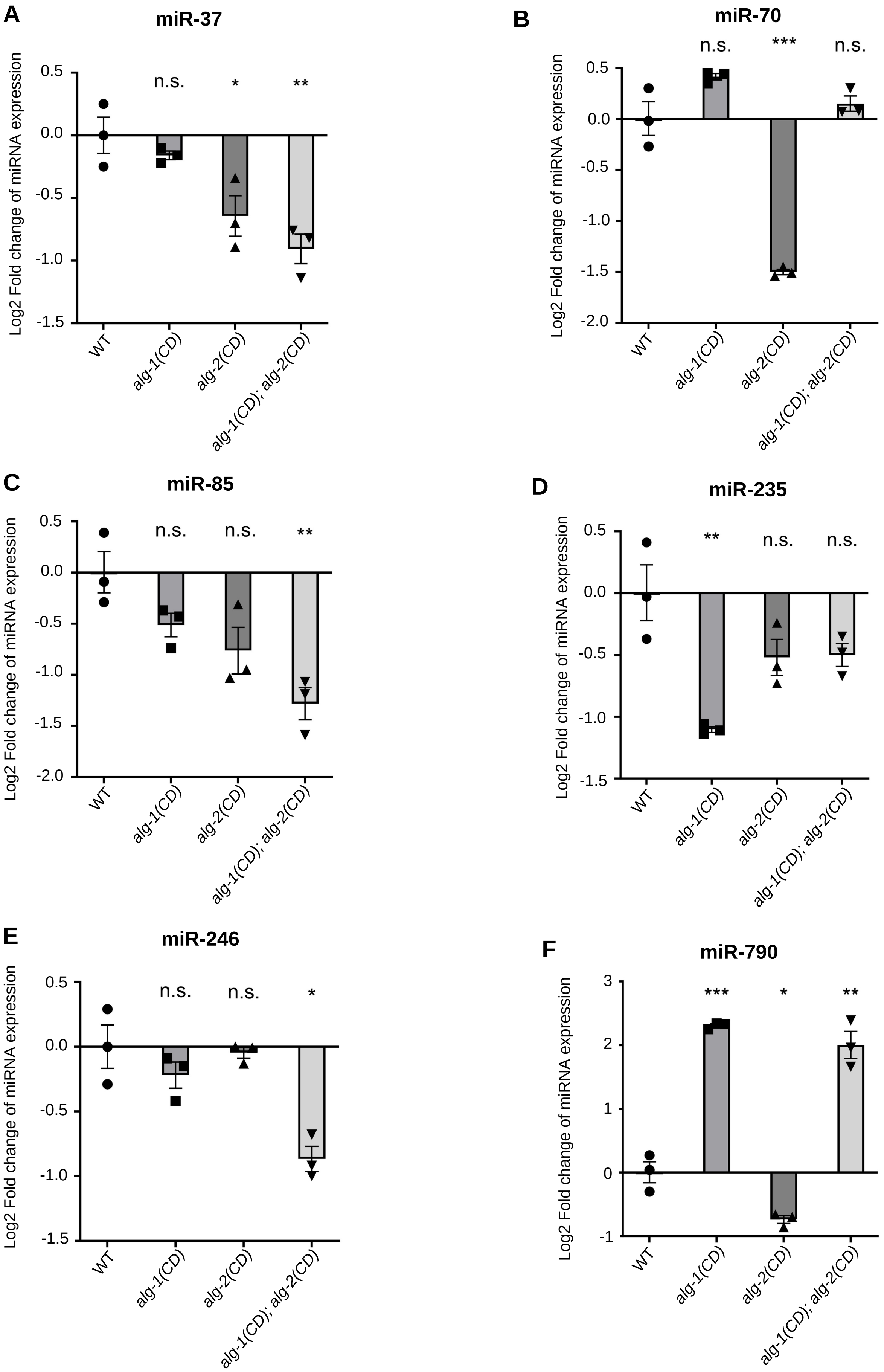
A
miR-37
Log2 Fold change of miRNA expression
0.5
0.0
-0.5
-1.0
-1.5
n.s.
*
**
WT
alg-1(CD)
alg-2(CD)
alg-1(CD); alg-2(CD)
B
miR-70
Log2 Fold change of miRNA expression
0.5
0.0
-0.5
-1.0
-1.5
-2.0
n.s.
***
n.s.
WT
alg-1(CD)
alg-2(CD)
alg-1(CD); alg-2(CD)
C
miR-85
Log2 Fold change of miRNA expression
0.5
0.0
-0.5
-1.0
-1.5
-2.0
n.s.
n.s.
**
WT
alg-1(CD)
alg-2(CD)
alg-1(CD); alg-2(CD)
D
miR-235
Log2 Fold change of miRNA expression
0.5
0.0
-0.5
-1.0
-1.5
**
n.s.
n.s.
WT
alg-1(CD)
alg-2(CD)
alg-1(CD); alg-2(CD)
E
miR-246
Log2 Fold change of miRNA expression
0.5
0.0
-0.5
-1.0
-1.5
n.s.
n.s.
*
WT
alg-1(CD)
alg-2(CD)
alg-1(CD); alg-2(CD)
F
miR-790
Log2 Fold change of miRNA expression
3
2
1
0
-1
***
*
**
WT
alg-1(CD)
alg-2(CD)
alg-1(CD); alg-2(CD)

**Figure S7**

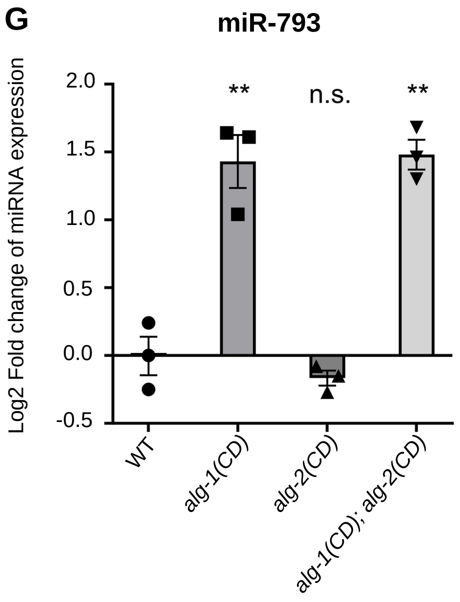


### Figure S7

**Figure S7. Quantification of miRNA expression by means of Taqman RT-qPCR. (A-G)** Levels of different miRNAs, detected from total RNA extracted from worms (N=3). U18 and Sn2841 have been used as internal controls. Mean along with S.E.M and individual data points for each miRNA tested, for each genotype and replicate are indicated. The *P*- value was obtained by unpaired two tailed t-test (**p*<0.05, ***p*<0.01, ****p*<0.001).

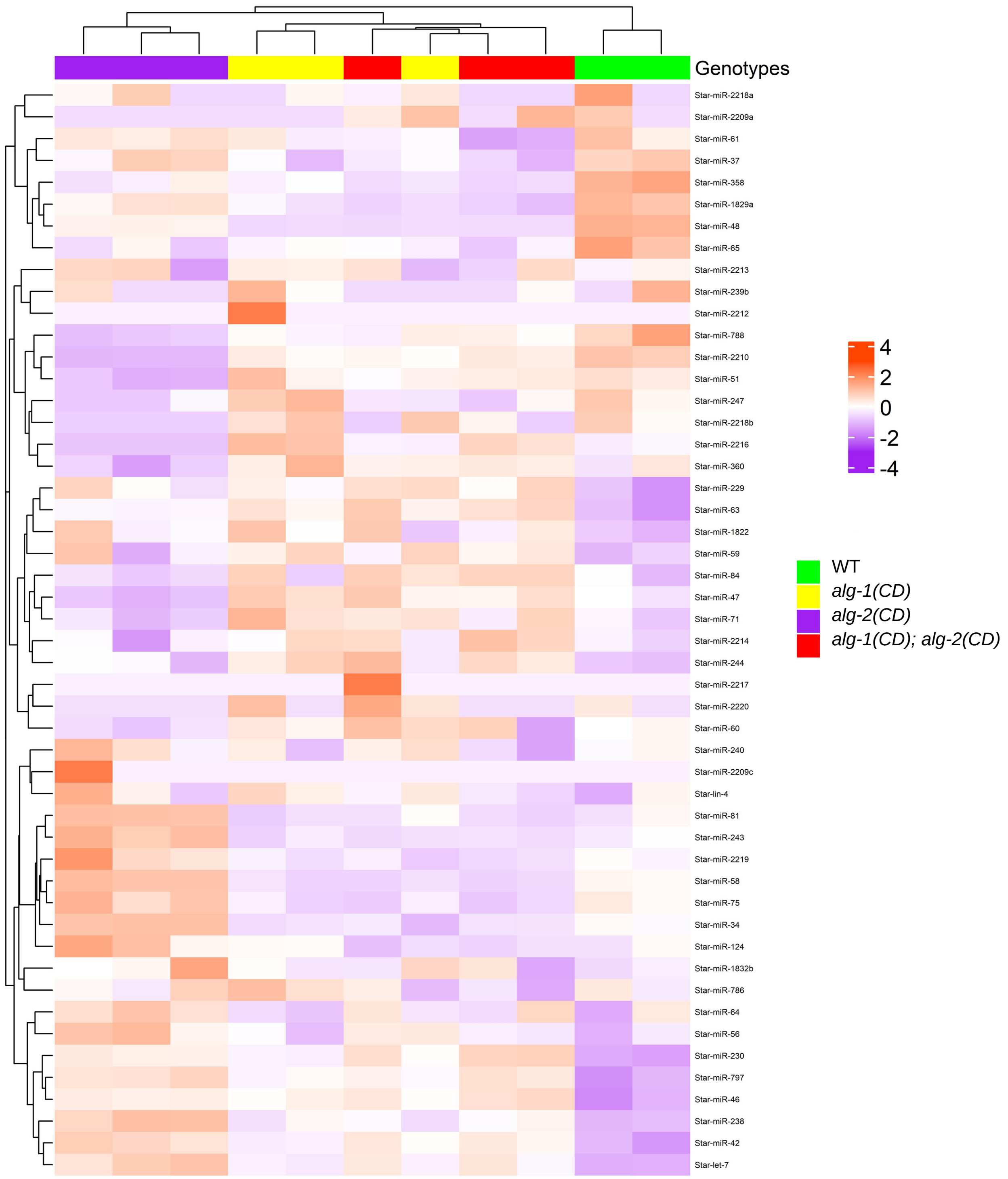
Genotypes
Star-miR-2218a
Star-miR-2209a
Star-miR-61
Star-miR-37
Star-miR-358
Star-miR-1829a
Star-miR-48
Star-miR-65
Star-miR-2213
Star-miR-239b
Star-miR-2212
Star-miR-788
Star-miR-2210
Star-miR-51
Star-miR-247
Star-miR-2218b
Star-miR-2216
Star-miR-360
Star-miR-229
Star-miR-63
Star-miR-1822
Star-miR-59
Star-miR-84
Star-miR-47
Star-miR-71
Star-miR-2214
Star-miR-244
Star-miR-2217
Star-miR-2220
Star-miR-60
Star-miR-240
Star-miR-2209c
Star-lin-4
Star-miR-81
Star-miR-243
Star-miR-2219
Star-miR-58
Star-miR-75
Star-miR-34
Star-miR-124
Star-miR-1832b
Star-miR-786
Star-miR-64
Star-miR-56
Star-miR-230
Star-miR-797
Star-miR-46
Star-miR-238
Star-miR-42
Star-let-7
4
2
0
-2
-4
WT
alg-1(CD)
alg-2(CD)
alg-1(CD); alg-2(CD)

**Figure S8**

**Figure S8. The effect of ALG-1 and 2 catalytic activities on miRNA passenger strand level.** Heatmap depicting the overall abundance of miRNA passenger strands in young adults in mentioned genotype. The tile color represents the number of standard deviations above (Red) or below (Blue) the mean expression of a miRNA (Known as Z-Score). Agglomerative hierarchical clustering based on Euclidean distance is performed on the samples (columns) and miRNA (rows), which is represented by their respective dendrograms.

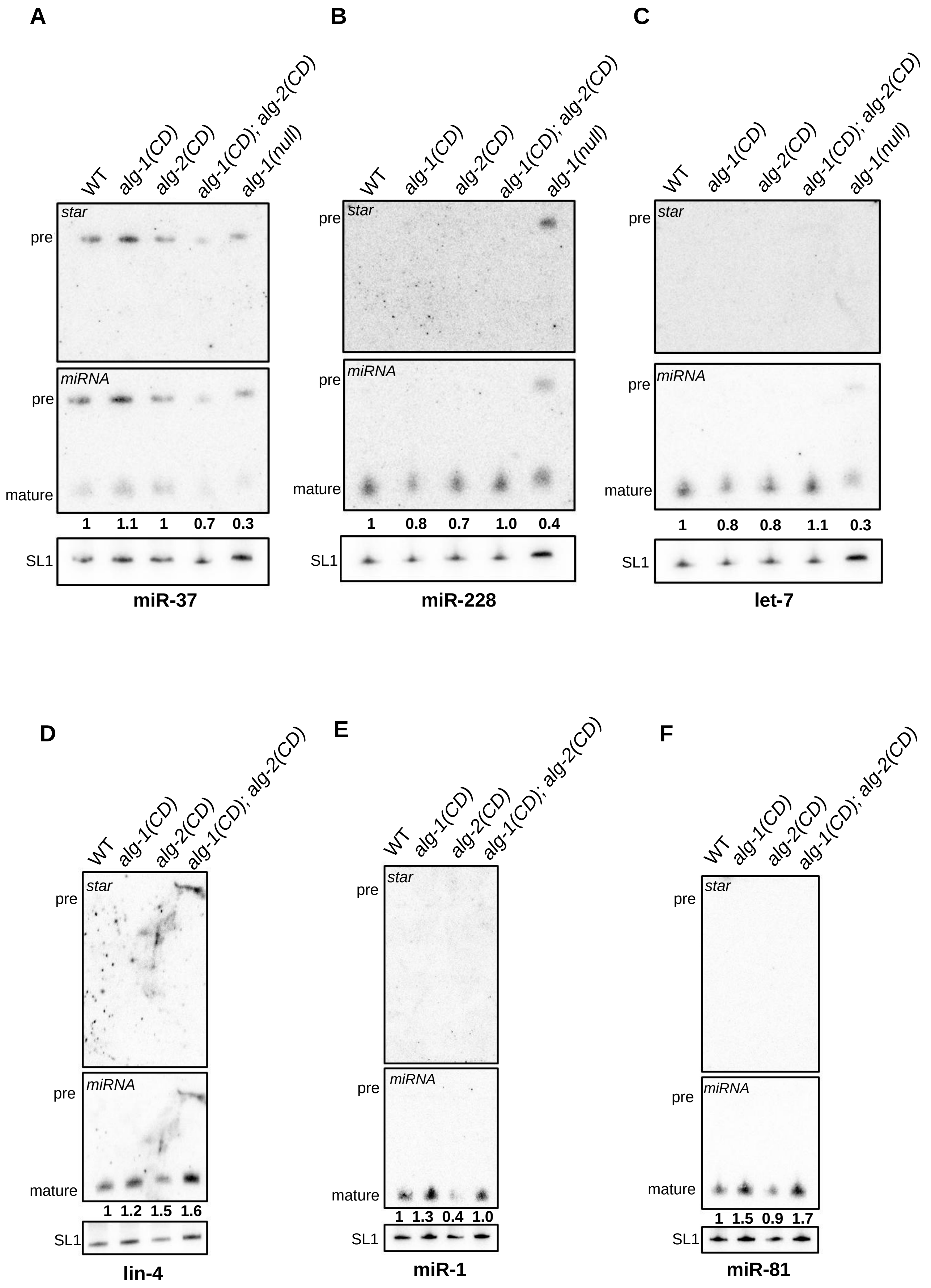
A
B
C
WT
alg-1(CD)
alg-2(CD)
alg-1(CD); alg-2(CD)
alg-1(null)
star
miRNA
pre
mature
1 1.1 1 0.7 0.3
SL1
miR-37
1 0.8 0.7 1.0 0.4
miR-228
1 0.8 0.8 1.1 0.3
let-7
D
E
F
1 1.2 1.5 1.6
lin-4
1 1.3 0.4 1.0
miR-1
1 1.5 0.9 1.7
miR-81

**Figure S9**

**Figure S9. Altering the catalytic activity of ALG-1 and ALG-2 does not influence the production of miRNAs. (A-F)** Both guide and star strands miRNAs were detected in young adults after total RNA extraction, followed by Northern blot. Complementary probes were used to detect specific miRNA strands (Table S5). *alg-1(null)* is used as a positive control **(A-C)** for the detection of precursor form of miRNA. SL1 RNA was probed and used as loading control. The relative intensity for mature miRNA(s) for each miRNA, probed for each genotype has been quantified against wild type.

Phenotype on specific RNAi

| Temp. | Genotype | RNAi | Fertility (%) (Mean±SEM) | Lethality (%) (Mean±SEM) | Sterility (%) (Mean±SEM) | Total (*n*) |
|---|---|---|---|---|---|---|
| 25°C | *wt* | 2XT7 | 100±0 | 0±0 | 0±0 | 635 |
| | | alg-1 | 94.68±0.37 | 5.32±0.37 | 0±0 | 588 |
| | | alg-2 | 99.85±0.15 | 0.15±0.15 | 0±0 | 681 |
| | *alg-1(CD)* | 2XT7 | 100±0 | 0±0 | 0±0 | 699 |
| | | alg-2 | 99.71±0.29 | 0.29±0.29 | 0±0 | 648 |
| | *alg-2(CD)* | 2XT7 | 99.85±0.15 | 0.15±0.15 | 0±0 | 542 |
| | | alg-1 | 94.96±0.28 | 5.04±0.28 | 0±0 | 614 |
| | *alg-1(null)* | 2XT7 | 92.61±3.34 | 7.39±3.34 | 0±0 | 594 |
| | | alg-2 | 0±0 | 5.3±1.82 | 94.7±1.82 | 508 |
| | *alg-2(null)* | 2XT7 | 99.42±0.58 | 0.58±0.58 | 0±0 | 344 |
| | | alg-1 | 0±0 | 8.33±0.67 | 91.67±0.67 | 391 |
| 15°C | *wt* | 2XT7 | 99.48±0.35 | 0.52±0.35 | 0±0 | 762 |
| | | alg-1 | 97.3±0.47 | 2.7±0.47 | 0±0 | 783 |
| | | alg-2 | 100±0 | 0±0 | 0±0 | 742 |
| | *alg-1(CD)* | 2XT7 | 100±0 | 0±0 | 0±0 | 595 |
| | | alg-2 | 99.43±0.12 | 0.57±0.12 | 0±0 | 687 |
| | *alg-2(CD)* | 2XT7 | 100±0 | 0±0 | 0±0 | 406 |
| | | alg-1 | 98.98±0.24 | 1.02±0.24 | 0±0 | 403 |
| | *alg-1(null)* | 2XT7 | 88.11±1.76 | 11.89±1.76 | 0±0 | 600 |
| | | alg-2 | 0±0 | 12.61±0.98 | 87.39±0.98 | 767 |
| | *alg-2(null)* | 2XT7 | 99.53±0.01 | 0.47±0.01 | 0±0 | 642 |
| | | alg-1 | 0±0 | 15.51±3.19 | 84.49±3.19 | 623 |

*alg-1(null)* worms are sick in general and at 25°C they are affected more.

#### Figure S10

**Figure S10. Phenotypic analysis of catalytic dead ALG-1/2 mutants exposed to RNAi mediated knockdown of *alg-1/2*.** The phenotypes, observed in wt and catalytic dead argonaute mutants, exposed to RNAi food targeting *alg-1/2*. Control RNAi (2XT7) acts as a negative control. The result is compiled from experimental replicates (N=3) and the mean and S.E.M. values have been determined for each phenotype for each genotype, exposed to specific RNAi food.

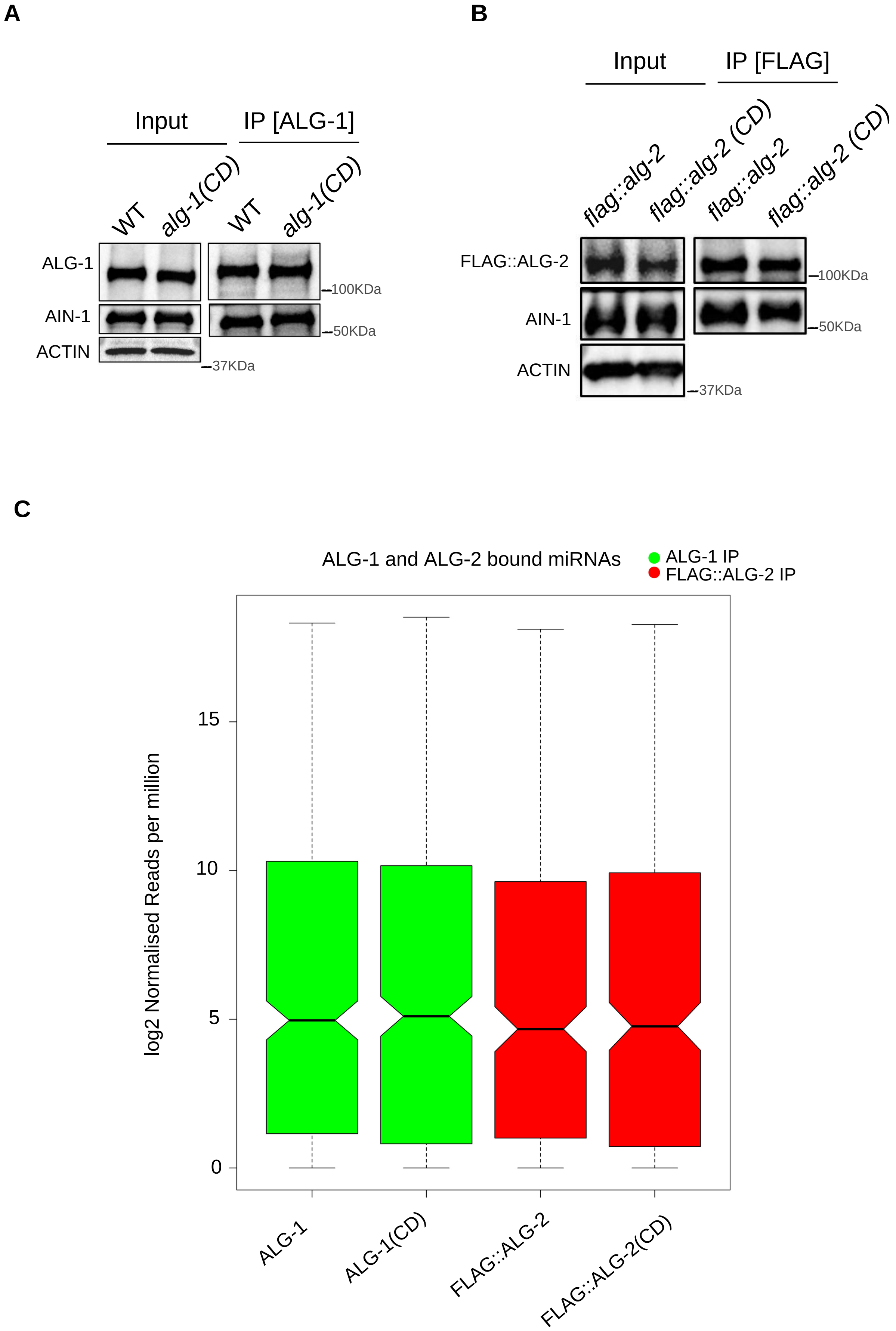
A
Input
IP [ALG-1]
WT
alg-1(CD)
WT
alg-1(CD)
ALG-1
AIN-1
ACTIN
100KDa
50KDa
37KDa
B
Input
IP [FLAG]
flag::alg-2
flag::alg-2 (CD)
flag::alg-2
flag::alg-2 (CD)
FLAG::ALG-2
AIN-1
ACTIN
100KDa
50KDa
37KDa
C
ALG-1 and ALG-2 bound miRNAs
ALG-1 IP
FLAG::ALG-2 IP
log2 Normalised Reads per million
0
5
10
15
ALG-1
ALG-1(CD)
FLAG::ALG-2
FLAG::ALG-2(CD)

**Figure S11**

**Figure S11. The loss of ALG-1 and ALG-2 catalytic activity does not affect their interaction with GW182 protein AIN-1 and miRNAs. (A)** Western blot of AIN-1 after ALG-1 immunoprecipitation from total protein extracts prepared with *alg-1(CD)* and wild-type embryos. **(B)** Western blot of AIN-1 after Flag::ALG-2 immunoprecipitation from total protein extracts prepared with *flag::alg-2(CD)* and *flag::alg-2* embryos. **(A-B)** Input is shown in the left panels and Actin is shown as a loading control. The images are representative of three replicates. **(C)** Boxplot of log2 Normalised Reads per million of detected guide miRNAs in ALG-1 (green) and FLAG::ALG-2 (red) immunoprecipitated complexes. There is no difference in the overall abundance of miRNAs associated to ALG-1(CD) or ALG-2(CD) when compared to the abundance of miRNAs associated to their respective wild-type counterparts.

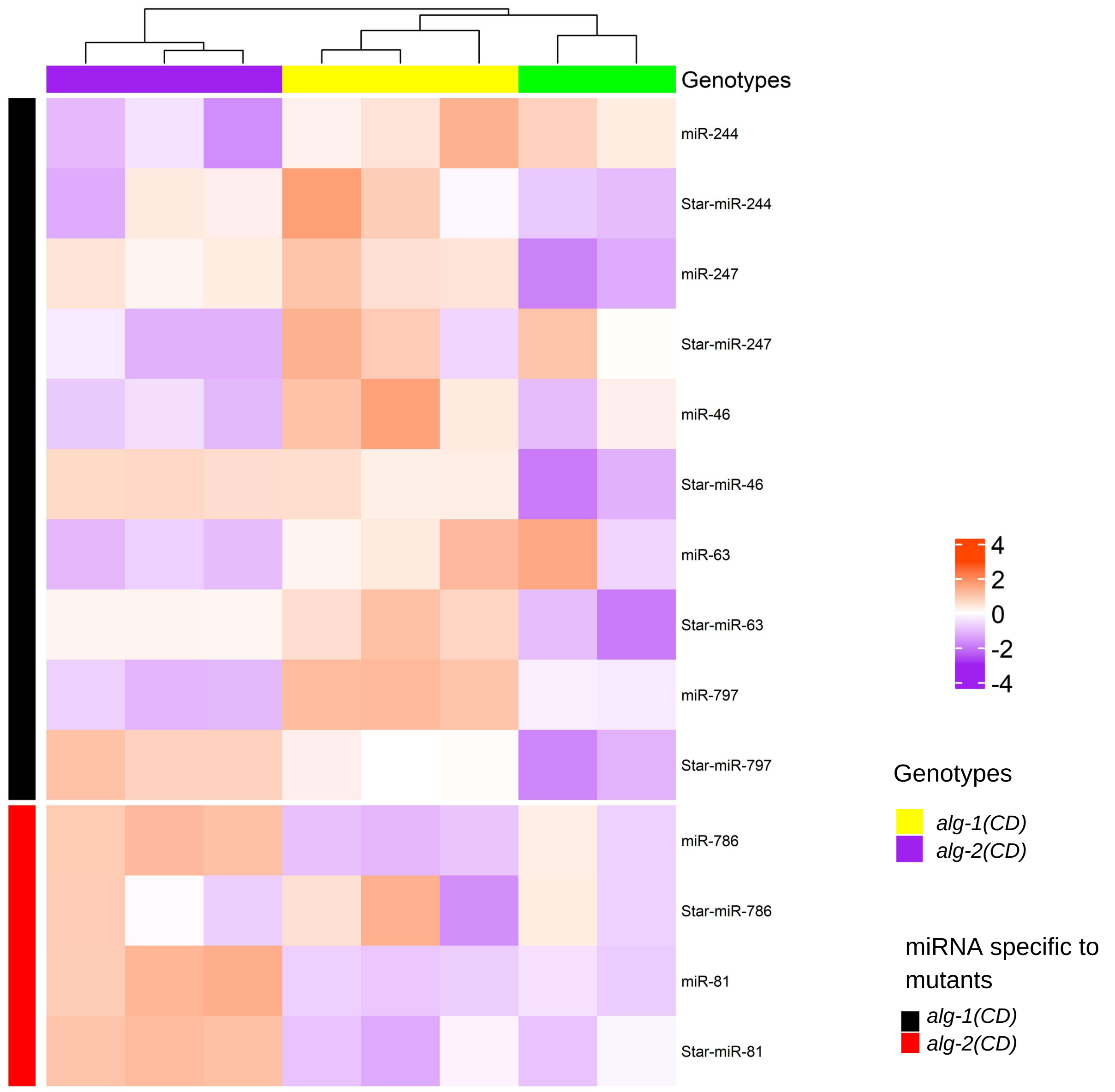


#### Figure S12

**Figure S12. Effect of the slicing activity of ALG-1 and ALG-2 on guide and star miRNA pairs.** Heatmap depicting abundance of guide and star strand miRNAs (pair) found to be significantly upregulated (guide and/or star strands) in either *alg-1(CD)* or *alg-2(CD)* mutant animals when compared to wild-type animals. The heatmap is split into two blocks based on the miRNAs and mutant specific upregulation and clustered by samples to verify for similarity among samples and genotypes for these specific guide and star pairs. The tile color represents the number of standard deviations above (Red) or below (Blue) the mean expression of a miRNA (Known as Z-Score). Agglomerative hierarchical clustering based on Euclidean distance is performed on the samples (columns) and miRNA (rows), which is represented by their respective dendrograms.

#### SUPPLEMENTARY TABLES (EXCEL FILE)

**Table S1:** Oligonucleotides used for this study
**Table S2-S4:** The details of CRISPR mutants, generated in this study
**Table S5:** Strains used in this study
**Table S6:** Probes used for Northern Blot
**Table S7:** Young adult samples used for sequencing
**Table S8:** Embryos samples used for sequencing
**Table S9:** Normalised Reads Per Million of all young adult samples
**Table S10:** Normalised Reads Per Million of all embryos samples
**Table S11:** Over Representation Analysis Of Significant Downregulated miRNA Targets
